# Helicase-assisted continuous editing for programmable mutagenesis of endogenous genomes

**DOI:** 10.1101/2024.02.01.577593

**Authors:** Xi Dawn Chen, Zeyu Chen, George Wythes, Yifan Zhang, Benno C. Orr, Gary Sun, Ka Thao, Mounica Vallurupalli, Jing Sun, Mehdi Borji, Emre Tkacik, Haiqi Chen, Bradley E. Bernstein, Fei Chen

**Author notes:** These authors contributed equally (Xi Dawn Chen, Zeyu Chen).

## Abstract

A major challenge in human genomics is to decipher the context specific relationship of sequence to function. However, existing tools for locus specific hypermutation and evolution in the native genome context are limited. Here we present a novel programmable platform for long-range, locus-specific hypermutation called helicase-assisted continuous editing (HACE). HACE leverages CRISPR-Cas9 to target a processive helicase-deaminase fusion that incurs mutations across large (>1000 bp) genomic intervals. We applied HACE to identify mutations in MEK1 that confer kinase inhibitor resistance, to dissect the impact of individual variants in SF3B1-dependent mis-splicing, and to evaluate noncoding variants in a stimulation-dependent immune enhancer of CD69. HACE provides a powerful tool for investigating coding and noncoding variants, uncovering combinatorial sequence-to-function relationships, and evolving new biological functions.

**One Sentence Summary:** We developed a tool for continuous, long-range, targeted diversification of endogenous mammalian genomes and used it to explore the function of genetic variants in both coding and non-coding regions.

## Main Text

A fundamental challenge of genomics is to chart the impact of the three billion bases in the human genome on protein function and gene regulation. Therefore, a critical goal is to develop strategies for mutagenizing genomic sequences systematically and at high throughput. In particular, saturation mutagenesis of single genomic loci could emulate the natural evolution process to reveal sequence-structure relationships, gain-of-function, and loss-of-function phenotypes. By performing such mutagenesis and selection in a stepwise and/or continuous process, this evolutionary process could be directed to generate enhanced protein functions, gene expression, or cell fitness.

However, targeted mutagenesis in the endogenous mammalian genome remains difficult for three primary reasons. First, many existing tools require exogenous overexpression of the gene of interest on a plasmid or vector (e.g., deep mutational scanning^1^, VEGAS^2^). This is sensitive to gene dosage and cannot be used to evolve noncoding regions in their native chromatin contexts. Second, some tools (e.g., TRACE^3^, TRIDENT^4^) require integrating exogenous sequences into the genome, which leads to experimental complexity and constraints throughput. Third, existing tools targeting the endogenous genome are either non-specific (e.g., alkylators^5^ that introduce genome-wide mutations) or confined to narrow genomic windows (e.g., CRISPR base-editors^6,7^, CRISPR-X^8^, or TAM^9^). Whereas CRISPR base-editor screens have been used to interrogate protein function and regulatory elements, they are limited in the base positions that can be targeted with high efficiency and can lead to artificial variants linkage due to the base editor mutating multiple bases in the editing window^10^.

Given the existing challenges in performing targeted and continuous mutagenesis in mammalian cells, we sought to develop a new platform for mutagenizing endogenous loci in their native chromatin contexts. We envisioned a tool with several advantageous properties, including (1) a long mutagenesis range (>200 bp); (2) the capacity to incur multiple, potentially interacting mutations across a region of interest; (3) a continuous and potentially tunable mutation rate for sampling variant space and exploring fitness landscape changes; and (4) a generalizable technical framework to target genomic loci of interest individually and in combination.

Here, to design such a tool, we introduce Helicase Assisted Continuous Editing (HACE), which combines long-range editing of entire loci with the advantages in sequence programmability inherent to CRISPR gene editing tools. HACE utilizes CRISPR-Cas9 to direct the loading of a helicase-deaminase fusion for targeted hypermutation of the downstream genomic sequence (Fig. 1, A and B). We evaluated HACE prototypes incorporating diverse helicases, nickases, and deaminases, showing that they have tunable edit rates and ranges. We then applied HACE to functionally characterize both coding and non-coding genomic elements. In the coding space, we identified variants that lead to MEK1-inhibitor drug resistance and also identified variants in splicing-factor SF3B1 that lead to alternative 3′ splice-site usage. Turning to regulatory regions, we defined functional artificial variants in the enhancer regions of CD69 and pinpointed specific bases and motifs that mediate the impact of RUNX factors on CD69 regulation. HACE provides a novel strategy for saturating mutagenesis of coding genes and regulatory regions, identifying specific nucleotides that underlie genotype-phenotype associations, and evolving new cellular functions.

**Figure 1:**
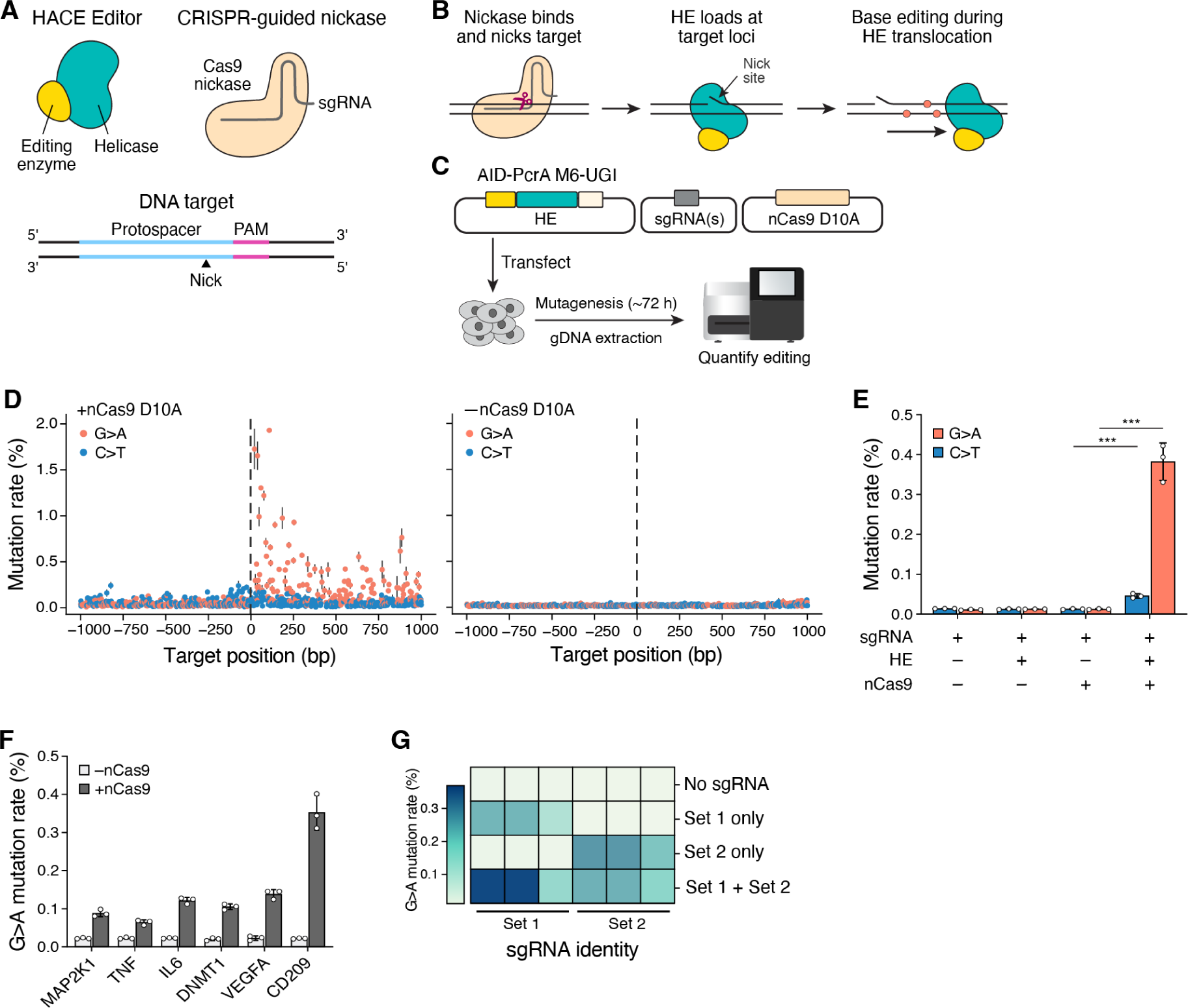
Overview of the helicase-assisted continuous editing (HACE) system. **(A)** Schematic of a HACE editor (HE), which is a fusion protein of helicase and base-editing enzymes (left). A CRISPR-guided nickase (Cas9 nickase) binds with a sgRNA to generate a single-stranded DNA nick at the target genomic DNA target (right). **(B)** Schematic model for HACE system. A CRISPR-guided nickase targets a specific genomic position to create a nick site. HACE editor (HE) loads to this site. As the helicase translocates along the DNA, the base editor introduces random point mutations (orange circles). **(C)** Schematic of HACE experimental workflow, involving co-transfection of a HE plasmid (PcrA helicase variant (PcrA M6) fused with a hyperactive mutant of activation-induced cytidine deaminase (AID*Δ) and uracil DNA glycosylase inhibitor (UGI)), a nSpCas9(D10A) plasmid, and sgRNA plasmid(s). Editing efficiency is assessed 72 hours post-transfection following genomic DNA extraction and amplicon sequencing. **(D)** Mutation rate per base across a ∼1-kb target region in the presence (left) and absence (right) of nCas9. The vertical dashed line shows the nick site. Data are mean of technical replicates (n= 3) ± s.d. **(E)** Mutation rate at the target loci (*HEK3*). The average mutation rate is calculated for both C>T and G>A modes for the region downstream of the nick site, excluding the sgRNA spacer region (Methods). Data are mean of technical replicates (n= 3) ± s.d. Significance is determined via unpaired two-tailed *t*-test between +/- nCas9 samples. \*\*\**P* < 0.001. **(F)** Mutation rate across genomic targets using loci-specific sgRNAs. The mutation rate is a sum of both C>T and G>A modes for the region downstream of the nick site, excluding the sgRNA spacer region. Data are mean of technical replicates (n= 3) ± s.d. **(G)** Average mutation rate for multiplex sgRNA targeting. Two sets of sgRNAs (three sgRNA per set, see Supplementary Table 1) are independently co-transfected with other HACE components. The mutation rate at each sgRNA target loci is depicted in the heatmap.

### HACE design and implementation

Helicases are highly processive enzymes that can traverse large genomic regions. Certain helicases, including those involved in DNA damage repair, can load and start unwinding DNA at nicks in the genome ^11^. We reasoned that such nick-directed helicases could be used for endogenous mutagenesis. We hypothesized that a helicase and deaminase enzyme fusion could generate multiple somatic edits across an extended genomic window. The fusion and its interval of hypermutation could be targeted to specific genomic regions using a nicking variant of SpCas9 (nCas9).

To test this, we constructed a HACE Editor (HE) protein by fusing a hyperactive mutant of activation-induced cytidine deaminase (AID*Δ) with a *Geobacillus stearothermophilus* PcrA helicase that was previously optimized for processivity (PcrA M6)^8,12^. We also appended a uracil DNA glycosylase inhibitor (UGI), which has been shown to facilitate C:G>T:A mutations^13^, at the 3′ end of the HE (Fig. 1C). To generate a targeted single-stranded nick, we used a SpCas9 nickase (nCas9; D10A) and corresponding single-guide RNA (sgRNA). After the nCas9 creates a nick, we reasoned that the HE would load at the nick, start unwinding the DNA, and generate random mutations in the process.

To this end, we co-transfected separate plasmids expressing HE, nCas9, and a sgRNA targeting the *HEK3* locus into HEK293FT cells. Cells were collected 72 hours after transfection, and editing rates were evaluated by amplicon sequencing. We observed directional editing from the nick-site in the presence of the HE, nCas9, and sgRNA (Fig. 1D). Importantly, we did not observe elevated mutation levels in cells transfected with only HE or only with nCas9 and sgRNA, suggesting that editing is driven by the HE and is guide-specific. Using the non-target strand of the Cas9 (strand that is not bound by the sgRNA) that runs from 3′ to 5′ as the frame of reference (Supplementary Fig. 1A), we observed a significant increase in mutation rates across a ∼1000 bp window downstream of the nick site, but not upstream, suggesting that editing occurs in a directional manner (Fig. 1D). We detected strand bias with G>A substitutions occurring at a higher rate than C>T substitutions, which likely reflects preferential repair of mismatches on the nicked strand ^13^. The direction of editing corresponds to helicase loading on the non-nicked strand and translocating in the 3′ to 5′ direction, which is consistent with known mechanisms of PcrA helicases ^14,15^. We quantified the editing rate downstream of the nick site, and we observed an average G>A mutation rate of 0.38% per base and an average C>T mutation rate of 0.046% per base (Fig. 1E, Methods), representing a significantly higher mutation rate than cells transfected with nCas9 or HE only (unpaired *t*-test, *P* < 0.001 for +/- nCas9 in both C>T and G>A groups). This also is a significantly elevated mutation rate as compared to the replication error rate of human cells ^16^. The rates of other transition and transversion mutation modes were comparable to the background, providing further support for the specificity and targeting of our fusion protein (Supplementary Fig. 1B).

To demonstrate that HACE can be targeted to diverse loci, we co-transfected the HE and nCas9 with different sgRNAs targeting regions that span a range of genomic contexts (tables S1 and S2). We observed elevated mutation rates in these loci only in the presence of sgRNA and nCas9, but not HE alone, suggesting that editing is guide and nCas9 dependent (Fig. 1F). In all locations, editing consistently occurs downstream of the sgRNA spacer (Supplementary Fig. 1C), confirming that editing is directional.

The modular architecture of HACE can also enable the simultaneous diversification of distinct genomic loci by coexpressing multiple sgRNAs. We co-expressed sets of three sgRNAs that target distant genomic loci. We observed elevated mutation rates in all target loci but not in untargeted control regions (Fig. 1G, Supplementary Fig. 1D). This suggests that HACE can couple hypermutation across distant genomic loci and thus enable multiplexing or evaluation of genetic interactions.

We also tested whether HACE could induce continuous nucleotide diversification over multiple cellular generations. We transfected the HACE constructs into HEK293FT cells and monitored the editing rate every 12 hours over 4 days. We observed a marked, continuous increase in mutation rate (Supplementary Fig. 1E), with the average number of mutations per contiguous Illumina sequencing read increasing over time points (Supplementary Fig. 1F). These data suggest that HACE can induce continuous nucleotide diversification across multiple cell generations, making it amenable to continuous evolution applications.

### HACE is flexible and programmable

We sought to optimize components for the HACE system to modulate its editing rate, range, and mutational modes. Broadly, the HACE system consists of four modular components: the helicase and editing enzyme that make up the HE, nCas9, and sgRNA that direct the fusion to target loci (Fig. 2A). We began by designing and testing HE constructs using six different monomeric helicases known to load at single-stranded nicks and shortlisted four for systematic profiling (Supplementary Fig. 2A, tables S3 and S4). We also tested two variants of nCas9, one that nicks the target strand (nCas9 D10A) and one that nicks the non-target strand (nCas9 H840A). We cloned constructs encoding different combinations of helicase and nCas9 variants, targeted the fusions to three different genomic loci (*HEK3*, *TNF*, and *IL6*) for examination of both mutation rate and editing range.

**Figure 2:**
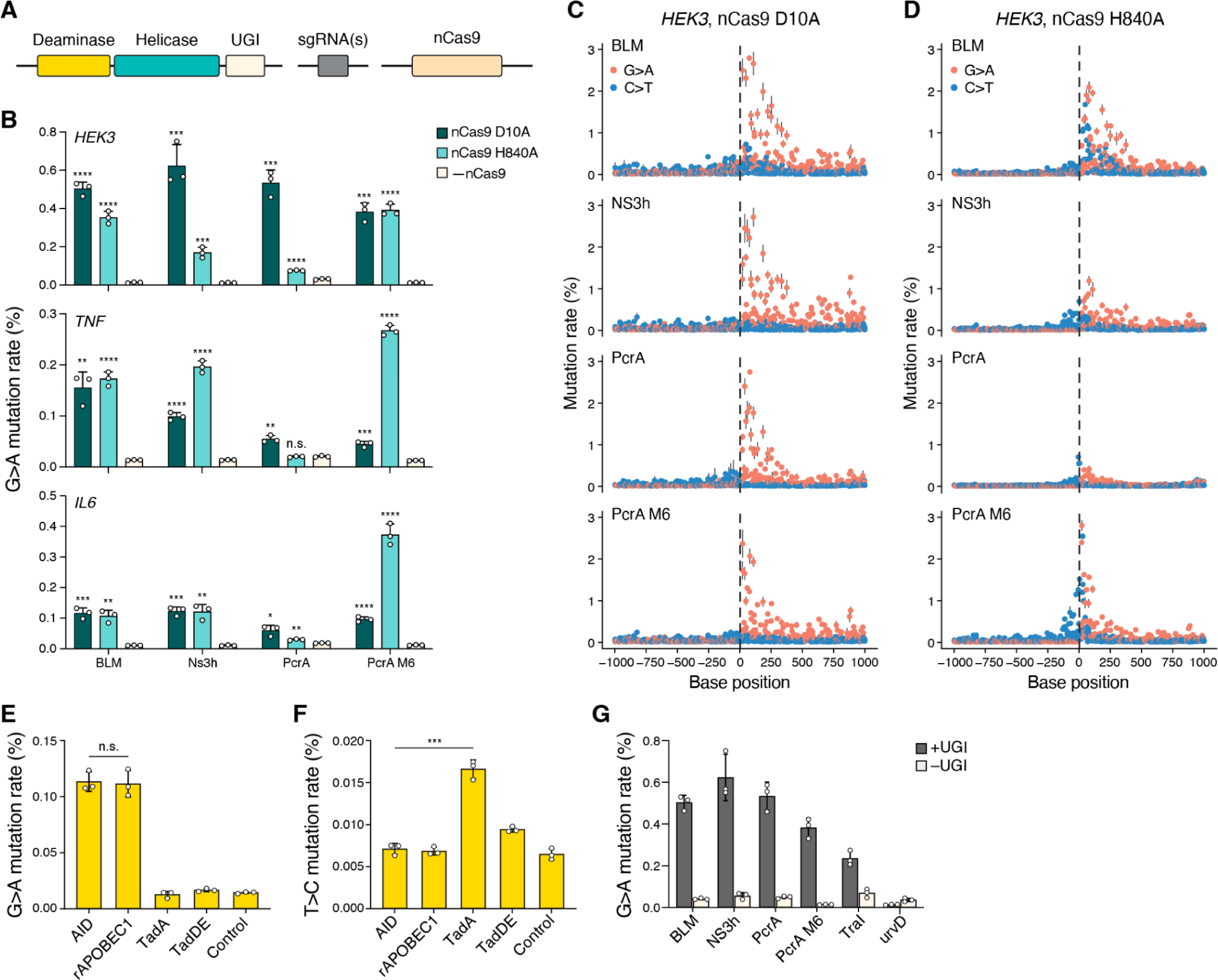
The HACE system is modular and flexible. **(A)** Modular components of the HACE system. Each component can be independently substituted to control for editing efficiency, mode, and range. **(B)** Mutation rate at the *HEK3*, *TNF*, and *IL6* loci for HEs with different helicase variants and the nCas9(D10A) or nCas9(H840A) nickase variant compared with (-) nCas9 condition. Significance is determined via unpaired two-tailed *t*-test between control and nCas9 samples with multiple-testing correction. n.s.: not significant. \**P* < 0.05. \*\**P* <0.01. \*\*\**P* < 0.001. \*\*\*\**P* < 0.0001. **(C)** Mutation rate per base across a ∼1-kb target region for HEs with different helicase variants and the nCas9(D10A) nickase variant. The vertical dashed line shows the nick site. Data are mean of technical replicates (n= 3) ± s.d. **(D)** Mutation rate per base across a ∼1-kb target region for HEs with different helicase variants and the nCas9(H840A) nickase variant. The vertical dashed line shows the nick site. Data are mean of technical replicates (n= 3) ± s.d. **(E)** Average G>A mutation rate at the *CD209* loci for HEs with different deaminase variants fused to the PcrA M6 helicase. Significance is determined via unpaired two-tailed t-test between AID and rAPOBEC1 groups. n.s.: not significant. **(F)** Average T>C mutation rate at the *CD209* loci for HEs with different deaminase variants fused to the PcrA-M6 helicase. Significance is determined via unpaired two-tailed *t*-test between AID and TadA groups. \*\*\**P* < 0.001. **(G)** Mutation rate at the HEK3 loci for HEs with different variants with and without UGI fusion. All data are mean of technical replicates (n= 3) ± s.d.

First, we tested HACE constructs with different helicase enzymes. We detected elevated mutation rates using either target-strand nickase (nCas9 D10A) or non-target-stand nickase (nCas9 H840A), with all constructs showing significant editing across the three loci for at least one nickase variant (Fig 2B). The preference of target and non-target strand nickase is loci-dependent, though the BLM helicase appears to prefer both nickase variants equally (Fig. 2B). We then characterized the editing range of different HE constructs. For each helicase, nickase, and loci combination, we transfected the respective HE, nCas9, and sgRNA combination into HEK293FT cells, harvested genomic DNA after three days, and then amplified a 1000 bp window by PCR for sequencing. The target-strand and non-target strand nickase had similar long-range editing performance (Fig. 2, C and D; Fig S2, B and C). Across a ∼1000bp window around the nick, we observed that the helicases still preferred to translocate in the downstream direction for the non-target strand nickase (nCas9 H840A), even though the nick is now on the opposite strand. We postulate that the nCas9 binding and DNA-sgRNA duplex might obstruct the helicase loading on the non-nicked single-stranded DNA.

To analyze the editing range, we calculated the local average mutation rate in 100 bp bins for each genomic loci (Supplementary Fig. 2D; Methods). We observed a decreasing mutation rate as a function of distance from the nick for all helicases profiled with both nickase variants. BLM, NS3h, and PcrA M6 helicases all demonstrated elevated editing (>10^-3^ G>A mutation rate per base) within 500 bp from the nick site. The mutation rate of PcrA M6 stabilized past 500 bp (at ∼10^-3^ G>A mutation rate per base), suggesting long, consistent, long-range editing up to 1000 bp away from the nick site. This range is an order of magnitude longer than previous Cas9-directed editing tools^8,9^.

Next, we sought to explore different base-editing enzymes for HACE. To identify the most efficient base editing enzyme and access diverse mutation modes, we engineered HEs fused with diverse deaminases, including (1) other cytosine deaminase enzymes that introduce C>T and G>A substitutions (rAPOBEC1^17^), (2) adenosine deaminase enzymes that introduce A>G and T>C substitutions (TadA-8e^18^), and (3) an engineered dual base editor that can perform both cytosine and adenine base editing (TadDE^19^). We tested these constructs in HEK293T cells and quantified mutation rates by amplicon sequencing (Fig. 2, E and F). We observed that rAPOBEC1 performs comparably to AID*Δ in introducing G>A base edits (unpaired *t*-test, *P* > 0.05). We found that TadA was able to induce T>C edits at a significantly higher rate than G>A editors (unpaired *t*-test, *P* < 0.001). On the other hand, the dual TadDE editor only induced minor levels of G>A and T>C editing. The deaminases rAPOBEC1 and TadA introduced mutations across diverse genomic loci (Supplementary Fig. 2E), demonstrating that HACE utilizing different deaminase fusions can introduce diverse programmable base editing modes.

Lastly, we found that the fusion of uracil glycosylase inhibitor (UGI) significantly elevated the editing levels for AID (Fig. 2G, unpaired *t*-test, *p* < 0.05 for all +/- UGI groups), consistent with reports from previous cytidine-base editor studies^13^. These results demonstrate that HACE editing rates can be tuned by varying the helicase and nickase variants used, making it suitable for diverse applications. Further engineering may leverage these insights to optimize the editing rate and range for specific applications.

### HACE is minimally perturbative in mammalian cells

While we found the HACE constructs to be well-tolerated in transfection experiments, we sought to quantify the effects of different HACE constructs on cell viability. To do so, we quantified cell viability using a luciferase-based ATP-assay (CellTiter-Glo) across various helicase constructs - both with and without deaminase - along with a loci-targeting sgRNA and nCas9 (Supplementary Fig. 3A). We found that HEs constructed with BLM and PcrA helicases did not result in a significant decrease in cell viability (unpaired *t*-test, p > 0.05 for each group).

However, AID-NS3h-UGI leads to a decrease in cell viability (unpaired *t*-test, *p* < 0.05), which is possibly related to the toxicity of NS3h helicases since it also acts on RNA^20,21^. AID-PcrA M6-UGI also significantly decreases cell viability (unpaired *t*-test, *p* < 0.001), while PcrA M6 alone did not affect cell viability (unpaired *t*-test, p=0.118). These results suggest that helicases used for HACE are well tolerated for cell viability.

Lastly, we explored whether HACE generated elevated mutation rates in non-targeted parts of the genome. To do so, we performed whole exome sequencing of cells expressing different HE variants and AID overexpression at high coverage (average ∼1000x coverage across the exome). To increase our statistical power to detect elevated mutation rates, we binned the genome into 100 kb bins and calculated the editing rate of each bin, then compared the editing rate between HE variants and control cells (Supplementary Fig. 3B). Of the 16621 genomic bins, we observed that overexpression of AID alone generated the most significant off-target bins (16 bins, Fisher’s Exact Test). We detected 5 bins with elevated editing rates for AID-NS3h-UGI, 2 for AID-PcrA-UGI and AID-PcrA M6-UGI, respectively, and 1 for AID-BLM-UGI. We found that bins with elevated editing rates in AID-NS3h-UGI overlapped with bins identified from other conditions, indicating common off-target sites across helicases (all sites outlined in Supplementary Table 5). Our data suggests that there is minimal elevation of global mutation rates in the exome due to HACE, and that off-target editing is likely driven by AID overexpression rather than the HE itself.

### HACE enables the identification of *MEK1* inhibitor-resistance mutations

Having developed and characterized the HACE system, we next sought to apply HACE to perform functional mutagenesis in both coding and non-coding genome contexts. In the coding genome, we first applied HACE to screen for mutations within mitogen-activated protein kinase kinase 1 (MEK1 kinase, also known as MAP2K1) that promote resistance to small-molecule drug inhibition. MEK inhibitors target the MAPK/ERK pathway, which is aberrantly upregulated in one-third of all cancers^22^. Using HACE, we diversified exons of the *MEK1* gene in A375 cells, a melanoma line sensitive to MEK inhibition, for three days, then selected cells for resistance to two MEK1 inhibitors – selumetinib and trametinib (Fig. 3A). We targeted exons 2, 3, and 6, which contain previously identified mutation hotspots^23^. Since the mutagenesis range of HACE is long, we only needed to design one sgRNA per exon. We placed each exon-specific sgRNA ∼100 bp upstream of the exon within the intronic region (Fig. 3B, Supplementary Table 1). After drug selection, we recovered surviving mutants and identified *MEK1* mutations by targeted amplicon sequencing (Methods). By comparing allele frequencies between pre- and post-drug selection samples and identifying alleles that are significantly enriched post-drug selection, we identified three candidate mutations that conferred resistance to trametinib (G128D, G202E, and E203K) and two candidate mutations that conferred resistance to selumetinib (G128D and E203K) (Fig. 3C; Supplementary Fig. 4; Supplementary Table 6). Two of the mutations, G128D and E203K, conferred resistance to both selumetinib and trametinib, suggesting similar resistance mechanisms.

**Figure 3:**
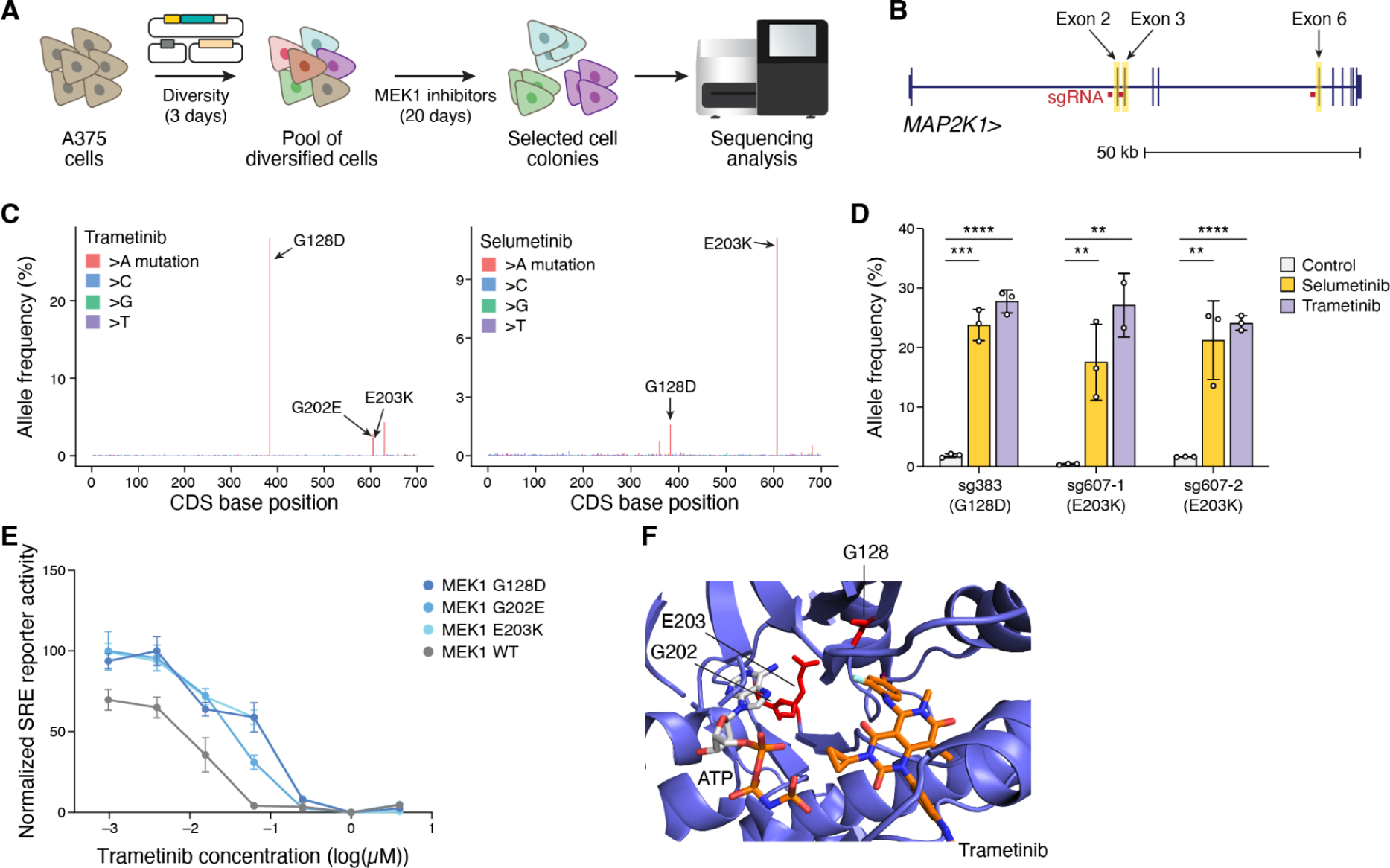
HACE enables the identification of *MEK1* inhibitor-resistance mutations in the endogenous genome. **(A)** Workflow of HACE MEK1 inhibitor resistance screen. A375 cells are transfected with HACE and diversified for 3 days. The genomic diversified cells are selected for 20 days. Genomes of resistant clones are harvested and sequenced by amplicon sequencing. **(B)** Location of sgRNAs for HACE screen. Exons 2, 3, and 6 (highlighted in yellow) are targeted for HACE diversification. Each exon-specific sgRNA (red bar) is placed ∼100bp upstream of the target exon. **(C)** Fold enrichment of *MEK1* cDNA sequence in trametinib-treated (left) and selumetinib-treated (right) samples. **(D)** Enrichment of mutations installed via base editing targeting G128D (sg383) and E203K (sg607-1/2) post trametinib or selumetinib treatment. Samples are sequenced 14 days post-selection by amplicon sequencing. Significance is determined via an unpaired two-tailed *t*-test between control and drug-selected samples. \**P* < 0.05. \*\**P* <0.01. \*\*\**P* < 0.001. \*\*\*\**P* < 0.0001. **(E)** MAPK–ERK signaling activity as measured by luciferase SRE reporter activity for G128D, G202E, and E203K mutants (mean ± s.e.m., n = 3 independent experiments). **(F)** Structure of MEK1 in complex with trametinib (PDB: 7JUR). MEK1 protein is depicted in blue, and residues that lead to drug resistance are annotated in red.

To validate top mutation candidates, we designed sgRNAs to introduce mutations individually into A375 cells using base editing, then selected edited cells with either selumetinib or trametinib for 14 days. We evaluated allele frequencies of introduced mutations pre- and post-selection by amplicon sequencing (Fig. 3D). We observed significant enrichment of G128D (sg383) and E203K (sg607-1 and sg607-2) post-selection with both inhibitors (sgRNAs in Supplementary Table 11). We could not introduce the 605G>A (G202E) mutation by base editing due to the artificial linkage in base editing between G202E and E203K. Therefore, we further validated candidates that conferred resistance to trametinib using a luciferase serum response element (SRE) reporter assay of MAPK-ERK signaling activity via exogenous overexpression of candidate MEK1 mutants (Fig. 3E, Supplementary Table 7). All three mutations individually increased trametinib resistance (IC_50_ = 68.0 nM, 46.1 nM, and 46.1 nM for G128D, G202E, and E203K, respectively, vs. 5.28 nM for wild-type). Structural analysis revealed that G128D is in the ligand-binding pocket. This mutation may function by inducing conformational changes of the binding pocket via steric interactions (Fig. 3F). On the other hand, E203K has been shown to cause constitutive MEK1 activation and downstream ERK phosphorylation, conferring gain-of-function ^24,25^. It is possible that the proximal G202E likely induces resistance through a mechanism similar to that of E203K. Overall, this demonstrates that HACE can effectively identify mutations conferring drug resistance while being less sensitive to the confounding effects of artificial genetic linkage. While previous screens have been performed in MEK1 kinase inhibitor resistance ^3,23^, this, to our knowledge, is the first mutagenesis-based resistance screen performed in the endogenous genome context without MEK1 overexpression.

### HACE enables the identification of variants in *SF3B1* that result in alternative 3′ branch point usage

Next, we applied HACE to explore the function of individual variants in splicing factor 3B subunit 1 (SF3B1) for splicing regulation. Mutations in RNA splicing factors occur in many cancer types and are especially prevalent in hematopoietic malignancies^26–28^. SF3B1 is the most frequently mutated splicing factor in cancer^29^. It is a member of the U2 small nuclear RNP (snRNP) complex and binds to the branch point nucleotide in the pre-catalytic spliceosome^30^. Pan-cancer analysis of SF3B1 mutations has identified hotspot mutations clustered within the C-terminal HEAT repeat domains 4-8 that display an alternative 3′ splice site (ss) usage signature^31,32^ (Fig. 4A). This mis-splicing occurs through the recognition of a different branch point sequence during 3′ss selection and results in global splicing changes associated with tumorigenesis. However, most known mutations identified from bioinformatic analysis of clinical samples have not been functionally validated for their effect on splicing. At the same time, clinically observed mutations only represent a small subset of the functional space of *SF3B1* variants. Identifying the mutations in SF3B1 that are functionally active in driving mis-splicing can improve understanding of splicing biology, predict future patient outcomes, and enable future therapeutic targeting.

**Figure 4:**
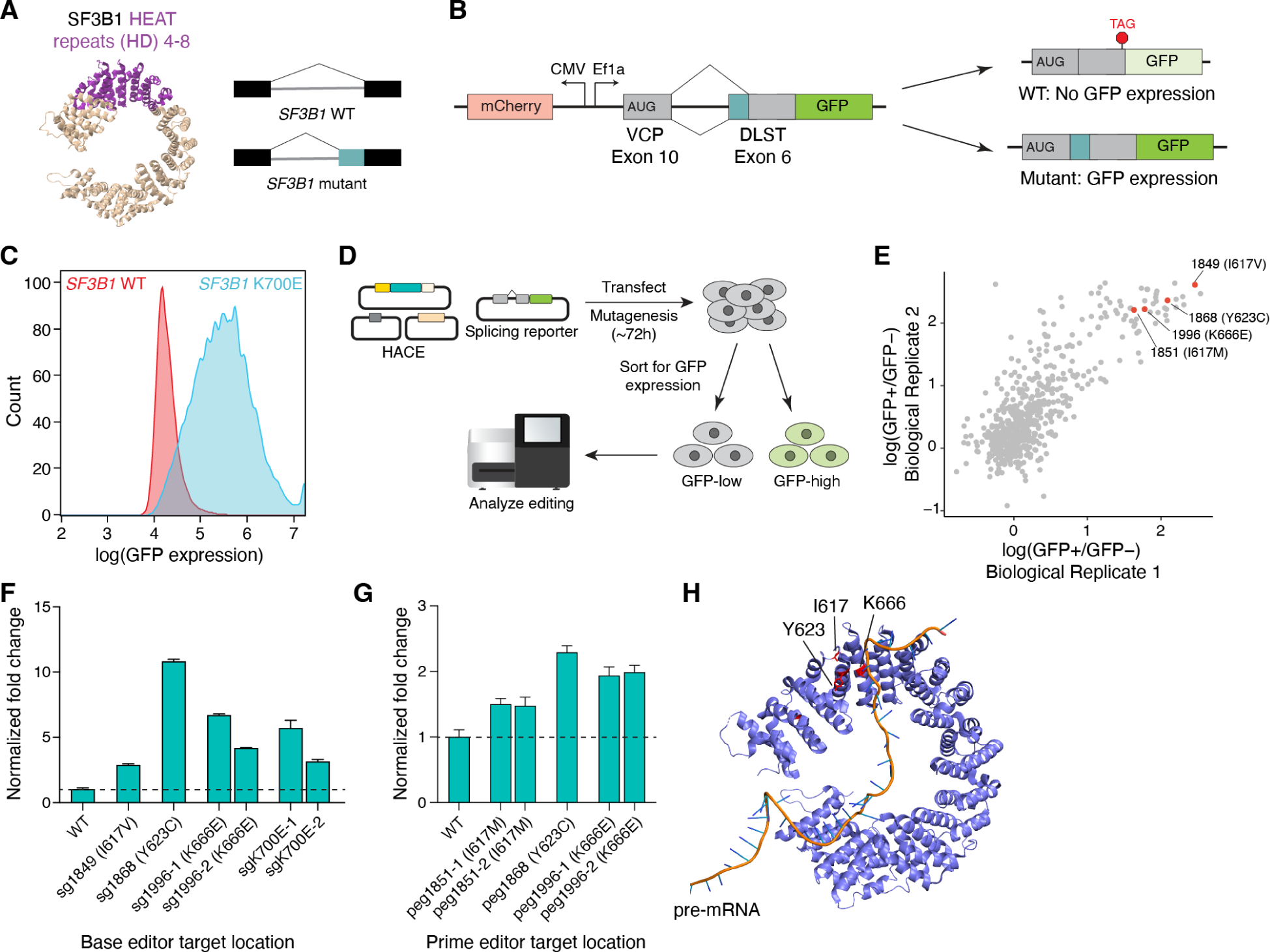
Identification of variants in SF3B1 that result in alternative 3′ branch point usage using HACE. **(A)** Structure of SF3B1 (right). HEAT repeats (HD) 4-8 are highlighted in purple (PDB: 6EN4). Differential splicing patterns can result from mutations in SF3B1 (left). **(B)** Schematic of the splicing reporter construct used for testing SF3B1-dependent splicing pattern. The plasmid reporter consists of a constitutively expressed mCherry, and a minigene splicing GFP reporter (VCP exon 10 fused with DLST exon 6 with a downstream GFP). Correct splicing will not generate GFP expression, while SF3B1-dependent altered splicing will lead to GFP expression. **(C)** Histogram of GFP signal measured by flow cytometry between isogenic K562 *SF3B1*^WT^ and *SF3B1*^K700E^ cells. Cells were gated for mCherry expression. **(D)** Schematic of *SF3B1* mutagenesis screen using HACE. HACE components and splicing reporter plasmids were co-transfected in HEK293FT cells. Mutagenesis was allowed to occur for 72h, and then cells were sorted for GFP expression. The editing rate for each sorted group was assessed following genomic DNA extraction and amplicon sequencing. **(E)** Fold enrichment of individual bases in the *SF3B1* cDNA sequence after selection across two biological replicates. Validated mutations are highlighted in red. **(F)** Normalized reporter activity fold change in mutations installed via base editing (mean ± s.d., n = 3). Significance is determined via an unpaired two-tailed *t*-test between control and edited samples. \*\*\**P* < 0.001 for all comparisons. The sgRNA sequences and their target bases are listed in Supplementary Table 11. **(G)** Normalized reporter activity fold change in mutations installed via prime editing (mean ± s.d., n = 3). Significance is determined via an unpaired two-tailed *t*-test between control and edited samples. \*\**P* < 0.01 for all comparisons. The epegRNA sequences and their target bases are listed in Supplementary Table 12. **(H)** Structure of SF3B1 (blue) in complex with pre-mRNA (orange). Validated mutations are shown in red. The structure was an overlay of PDB structures 6AHD and 5IFE.

To screen for mutations in SF3B1 that functionally lead to mis-splicing, we first sought to construct a minigene reporter that could distinguish between wild-type SF3B1 (*SF3B1*^WT^) and mutated SF3B1 splicing patterns. These patterns display alternative 3′ss usage characteristic of hematopoietic malignancies. First, we compared RNA-seq data from isogenic K562 cells containing either *SF3B1*^WT^ or mutant SF3B1 (*SF3B1*^K700E^, a mutation known to induce the alternative 3′ss phenotype). We shortlisted splicing events that were significantly differentially spliced between WT and mutant cells and constructed minigene reporters from two of the top sequences to test their ability to functionally distinguish between *SF3B1*^WT^ and *SF3B1*^K700E^-induced mis-splicing. To do so, we extracted the last 150bp of the endogenous intron and its downstream exon for each sequence and constructed a minigene by fusing it to a constant upstream exon and a downstream GFP reporter (Methods, Figure 4B, Supplementary Table 8). Our reporter design was such that correct splicing would result in a frameshift in the open reading frame of GFP, suppressing fluorescence, while mis-splicing would permit GFP expression. Each minigene-GFP sequence was cloned into a vector with constitutive mCherry expression, enabling quantitative assessment of mutation-dependent protein production through flow cytometry (Methods).

First, to validate our splicing reporter constructs, we transfected each construct into isogenic *SF3B1*^WT^ or *SF3B1*^K700E^ K562 cells and measured mutant-dependent protein expression by flow cytometry. Encouragingly, both reporters showed mutant-dependent specificity, showing elevated GFP expression in *SF3B1*^K700E^ cells compared to *SF3B1*^WT^ cells (fig S5A). We further confirm that alternative 3′ss usage drives the reporter expression using targeted RNA-seq (Supplementary Fig. 5B). We selected the reporter with the most significant mutant-dependent specificity derived from DLST Exon 6 to proceed with our *SF3B1* variant screen using HACE (Fig. 4C).

To perform the screen using HACE, we diversified exons 13-17 of the *SF3B1* gene in HEK293FT cells for three days. This region corresponds to HEAT repeat domains 4-8 of SF3B1, the region where known mutations for mis-splicing are enriched. Similar to the *MEK1* screen, we placed all exon-specific sgRNA within intronic regions covered by seven sgRNAs. To widen the genetic search space, we used HACE editors that can cover both C:G>T:A and A:T>G:C mutation modes (AID*Δ-PcrA M6-UGI and TadA-8e-PcrA M6-UGI). We transfected the minigene reporter into diversified cells, sorted the cells into two bins (GFP^–^ and GFP^+^) based on the GFP:mCherry ratio, and performed high-throughput sequencing for cells in each bin (Fig. 4D). Enriched mutations were identified by comparing the fold-change between GFP^–^ and GFP^+^ cells (Fig. 4E, Supplementary Table 9). We observed a high degree of replicate correlation for enriched variants between two independent biological replicates (Pearson’s ρ = 0.795). We also compared the highly enriched variants (>10-fold enrichment) to mutations observed in clinical datasets ^33^ and found nine variants that occurred at high frequency clinically (Supplementary Fig. 5C).

To validate the candidates that displayed the highest enrichment levels, we introduced the candidate mutations individually into HEK293FT cells using base editing in cells that were co-expressing the minigene reporter. We then quantified the fold change in GFP:mCherry ratio compared to unedited cells. We found that three of the mutations (I617V, Y623C, K666E) led to a significant increase in reporter fold-change (unpaired *t*-test, *p* < 0.001 for all base editing groups compared with control, Fig. 4F, Supplementary Fig. 5D). The reporter fold-change value was similar to fold change observed with base-editing installing K700E, a well-validated mutation affecting SF3B1 alternative 3′ss usage^34^. We also performed targeted amplicon sequencing for each cell population to validate editing at each target base (Supplementary Fig. 5E). We noted that two of the mutations, Y623C and K666E, have been observed in clinical datasets and are highly enriched in hematopoietic tumor samples^26,35^. K666E has been previously validated for its effect on alternative 3′ss usage^34,36^. We also observed an additional mutation, I617V/M, not previously observed in clinical datasets. We further validated these top candidate mutations via prime-editing and measured reporter fold change (Fig. 4G, Supplementary Fig. 5F, Supplementary Table 12). Despite the low editing efficiency, we observed significantly increased splicing reporter fold changes for the mutations as compared to WT cells (unpaired *t*-test, *p* < 0.01 for all prime editing groups compared with control). Overall, the editing rate for candidate mutations that affect SF3B1 alternative 3′ss usage across validation experiments correlates well with the minigene reporter fold change (Supplementary Fig. 5G).

Analysis of the protein structure of these mutations found that the mutations are all located at the edge of the HEAT repeat helices of the SF3B1 protein structure, which matches the pattern of hotspot mutations previously found from pan-cancer analysis (Fig. 4H). These mutations are located near the pre-mRNA-binding region and have been shown to disrupt the tertiary structure of the SF3B1 protein^37^. Taken together, HACE has helped us to identify clinically relevant mutations that result in SF3B1-dependent mis-splicing.

### Mutagenesis of noncoding regulatory elements uncovers functional bases and variants

We next leveraged HACE to identify functional bases and variants within non-coding gene regulatory elements. Resolving the individual bases and variants that underlie the activity of enhancers remains challenging as these and other genomic regulatory elements must ideally be examined in their native chromatin context. Although we and others have used CRISPR base editors to characterize enhancers^38–40^, such approaches require separate sgRNAs for each narrow target window and may be limited by the occurrence of PAM sites (only ∼30% of locations are targetable with NGG PAM sites). Furthermore, artificial linkage between variants due to correlated mutations in an editing window limits the ability of base editors to resolve the functions of single bases^41^.

We targeted HACE to an enhancer region that regulates *CD69*, a membrane-bound lectin receptor gene that contributes to immune cell tissue residency^42–45^. We designed three sgRNAs targeting the core region of the *CD69* enhancer (Supplementary Table 1). We infected K562 cells with these nCas9-sgRNAs and HE (AID-PcrA-M6-UGI) constructs. After 6 days, we stimulated the cells with PMA/ionomycin to induce CD69 expression and sorted cells based on CD69 surface expression (Methods). We assessed mutations by amplifying and sequencing the targeted region in CD69^low^ and CD69^high^ subsets (Fig. 5A and Supplementary Fig. 6A). The relative effect of each base edit was calculated based on its enrichment/depletion in CD69^high^ relative to CD69^low^ libraries (Methods). Multiple individual bases reduced CD69 activation, with most of them located in motifs of immune-related transcription factors (Fig. 5B, Supplementary Table 10). The base enrichment pattern was highly consistent across biological replicates(Pearson’s ρ = 0.845), confirming the robustness of our screen (Fig. 5C).

**Figure 5:**
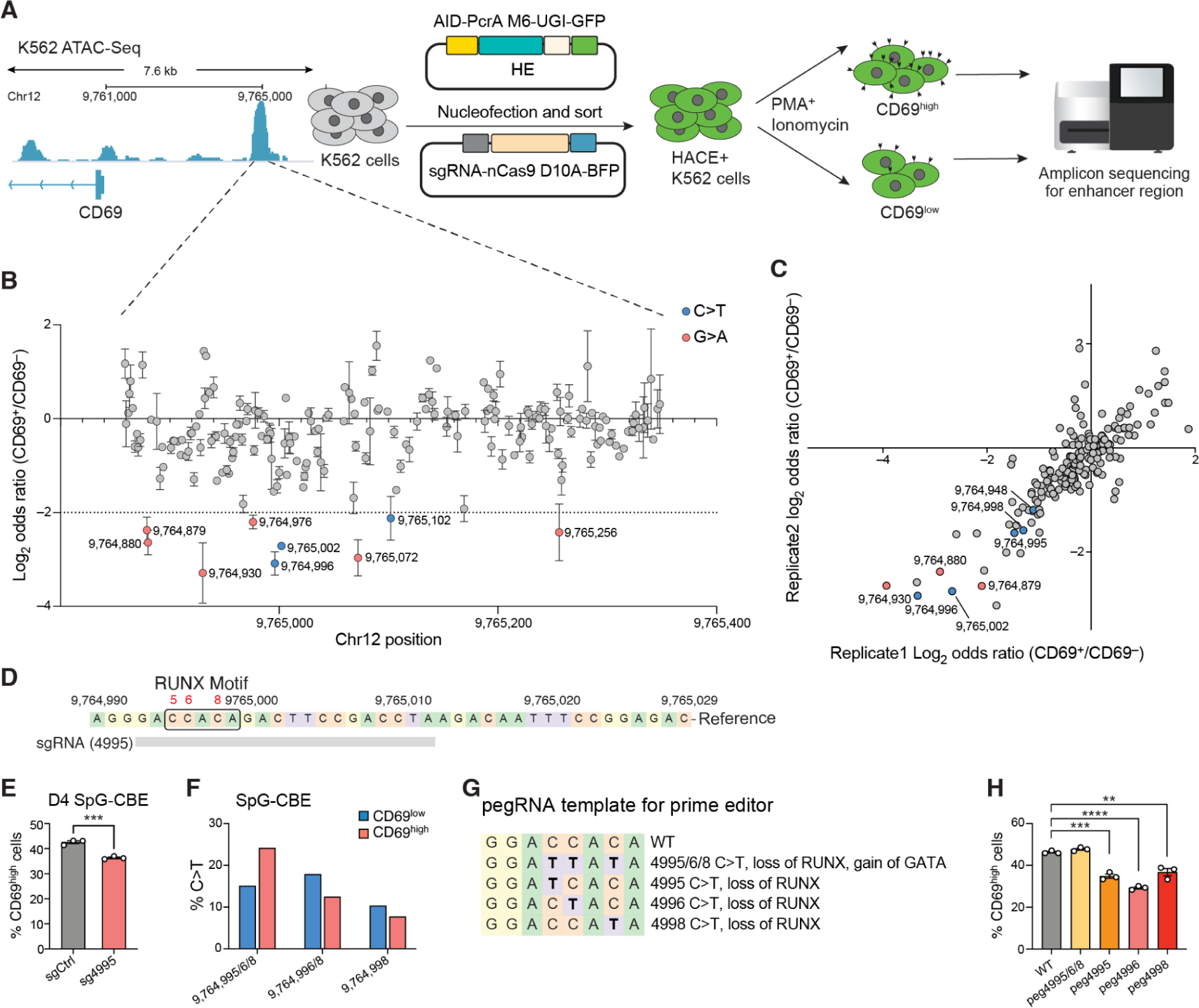
Single-base tuning of cis-regulatory elements via HACE identifies transcriptional regulation of CD69 by RUNX1/2. **(A)** Schematic of experimental workflow. The *CD69* enhancer region in K562 cells was identified using ATAC-seq data and targeted via HACE sgRNAs. HACE+ K562 cells were diversified for 6 days, then stimulated with PMA/ionomycin to induce CD69 expression. Cells are sorted into CD69^high^ and CD69^low^ populations, and the editing rate was profiled using amplicon sequencing. (B) Per base enrichment of C>T or G>A edits in CD69^high^ cells relative to CD69^low^ cells (Methods). The top most enriched C>T (blue) and G>A (red) variants in the CD69^low^ population are annotated. Each data point represents mean ± s.e.m. (n=2). **(C)** Fold enrichment of individual bases in the *CD69* enhancer region across 2 biological replicates. Validated bases are highlighted in blue (C>T) or red (G>A). **(D)** Sequence of chr12:9764990-9765029. RUNX motif boxed. A sgRNA (sg4995) with NG-PAM was used to target multiple cytosines in the RUNX motif. **(E)** Bar plot depicts the proportion of CD69^high^ cells in SpG-CBE-sgCtrl (gray) and SpG-CBE-sg4995 (red) after stimulation on day 4 post-transfection. Significance is determined via unpaired two-tailed *t*-test between groups (\*\*\**P* < 0.001). Data are from 3 independent experiments each with 3-4 technical replicates, mean ± s.e.m.. **(F)** Frequency of different incurred base edit combinations in sg4995-transfected K562 cells in CD69^low^ and CD69^high^ populations. **(G)** The pegRNA templates for single base dissection around chr12:9764992-9764999. The hypothesized changes in phenotype are annotated. **(H)** The proportion of CD69^high^ post-stimulation for cells edited with different pegRNAs on day 4 post transfection. Significance is determined via unpaired two-tailed *t*-test between WT and edited groups. \*\**P* <0.01. \*\*\**P* < 0.001. \*\*\*\**P* < 0.0001.

We validated the HACE screen by evaluating mutations that were located in three immune-related transcription factor motifs via base editing. We infected K562 cells with a base editor construct and sgRNAs designed to incur the corresponding edits, activated the cells, and analyzed CD69 expression by flow cytometry. We first confirmed that a C>T transition at base Chr12:9764948 (shortened as the last four digits “4948”, same nomenclature used below) significantly suppressed CD69 induction upon stimulation (Supplementary Fig. 6B). Our base editing tiling screen previously identified this artificial variant, which disrupts a GATA transcription factor motif^41^. We also validated G>A transitions at both bases 4879 and 4880 that suppress CD69 induction (Supplementary Fig. 6C). These variants lie within a predicted IRF/ETS or IRF/STAT transcription factor motif (“GAAAGGAA”), suggesting a role for the motif and cognate factors in regulating CD69 expression during stimulation.

Next, we focused on three closely adjacent screen hits: C>T transitions at positions 4995, 4996, and 4998. Motif analysis suggests these variants are located within the core motif region (“CCACA”) recognized by RUNX family transcription factors (Fig. 5D). RUNX1 and RUNX2 are both expressed in K562 cells and have previously been implicated in CD69 expression ^46,47^. Indeed, CD69 expression increased when we overexpressed either RUNX1 or RUNX2 (Supplementary Fig. 7A), and CD69 expression levels decreased when we knocked down RUNX1 or RUNX2 using shRNA (Supplementary Fig. 7B). These results support a role for a RUNX1/2 circuit in driving CD69 induction in K562 cells.

To validate these data, we designed a sgRNA targeting the 4995-4998 region and introduced C>T mutations using an NG-PAM cytidine base editor (Fig. 5D). We confirmed that the base edits reduced CD69 expression four days post-editing (Fig. 5E, Supplementary Fig. 7C). To evaluate the impact of different combinations of variants in this window, we sorted CD69^low^ and CD69^high^ populations and performed targeted amplicon sequencing. We found that edited alleles with a single mutation at position 4998 or paired mutations at positions 4996 / 4998 were enriched in the CD69^low^ population, consistent with an adverse effect on CD69 induction. This loss-of-function likely reflects the ablation of the coincident RUNX motif. However, edited alleles with C>T mutations at all three positions (4995/4996/4998) were enriched in the CD69^high^ population (Fig. 5F, Supplementary Fig. 7D-E). A likely explanation for this discordance is that the concurrent triple mutation not only disrupts the RUNX motif but also creates a GATA motif (“GATT”) at positions 4993-4996. GATA factors, including GATA1 and GATA2, are highly expressed in K562 cells and associated with transcriptional induction. These data suggest that de novo motif creation and GATA recruitment underlie the increased CD69 induction associated with the triple mutant allele (Fig. 5G).

Our findings support the potential of combinatorial base editing to create gain-of-function regulatory sequences but also highlight complications related to artificial linkage between adjacent variants. To definitively address this limitation, we used prime editing to introduce either individual or triple mutations at this locus (Supplementary Table 12). We again found that the triple mutation (4995/4996/4998) increased CD69 induction (Fig. 5H, Supplementary Fig. 7F), while the respective single mutations reduced induction. Amplicon sequencing of sorted cells further confirmed that the CD69^high^ population is enriched for the triple mutant allele but depleted for the respective single mutants (Supplementary Fig. 7, G and H). Hence, the application of HACE, followed by base and prime editing validation, revealed single nucleotide variants and combinatorial mutations capable of significantly modulating the activity of the CD69 immune enhancer through ablation or creation of transcription factor motifs. Our data demonstrate the ability of long-range HACE mutagenesis to dissect sequence determinants of regulatory element function.

## Discussion

Here, we developed HACE, a technology that enables programmable hypermutation of endogenous genomic loci in mammalian cells. HACE leverages CRISPR/Cas technology for targeted and multiplexable mutagenesis of the mammalian genome. HACE is unique both in its ability to distribute edits across large (>1000 bp) genomic regions and in its potential to evolve alleles over multiple cellular generations. We demonstrate HACE in forward genetic screens to identify coding mutations that confer resistance to clinical kinase inhibitors, to identify and validate clinically relevant mutations in the SF3B1 splicing factor that induce altered 3’ splice site selection, and to uncover artificial variants that alter the activity of an immune enhancer.

HACE provides key advantages over existing methods for mammalian mutagenesis (Supplementary Table 13). First, HACE solves the two long-standing limitations faced by conventional base editing screens: the requirement of an NGG PAM adjacent to the sgRNA recognition sequence and the occurrence of bystander mutations that can create artificial linkages and confound screening results. Instead of necessitating multiple sgRNAs to tile a target locus, the extended range of HACE confers mutations across a large genomic region using a single sgRNA, decreasing complexity and cost for pooled screens across multiple genomic targets. Second, the long editing range can uncover combinatorial effects and interactions between multiple distant mutations across a locus - a unique capability not afforded by existing base editor methods. HACE is also amenable to continuous mutagenesis over multiple cell generations. Compared to existing mammalian directed evolution strategies^48,49^, HACE offers an entirely new platform for laboratory evolution studies with the potential to chart fitness landscape changes, recapitulate evolutionary trajectories of biological phenotypes, or identify multi-step synergistic mutations that confer de novo protein or gene regulatory functions within the endogenous mammalian genome.

While powerful, our existing HACE system is subject to certain limitations that should be addressed through further engineering. One goal would be to increase mutation rates and range by optimizing modular components of the HACE system. For example, a wide diversity of helicases can be explored for improved efficiency, processivity and fidelity, and faster kinetics. Such efforts could leverage insights from the rational engineering of helicases for other applications and potentially allow for helicases with adjustable mutation ranges, which could help restrict mutations to relevant genomic windows (e.g., enhancers). Second, HACE is currently limited to transition mutation modes due to the use of cytidine and adenosine deaminases but might be expanded by incorporating emerging base-editing enzymes^50–52^. Lastly, further computational tools for optimal guide design and experimental approaches for HACE expression regulation may enable long-term HACE editing in a variety of cell types.

Overall, HACE represents the first example of continuous, long-range, programmable diversification of endogenous mammalian genomes. We envision HACE will significantly expand the functional genomics toolbox and unlock new molecular-scale insights toward building sequence-function maps of both coding and noncoding genomes.

## Acknowledgments

We thank the members of the Chen and Bernstein labs for helpful discussions. We would also like to thank Aarti Krishnan, Lindsey Guan, and Alexa Guan for their support and helpful discussions. We also thank Andrew Patentreger, AJ Masse, and Natan Pirete at the Broad Institute Flow Cytometry Core Facility for their experiment support.

## Authors’ contributions

F.C., G.W., and X.D.C. conceived the study. X.D.C., Z.C., B.O., Y.Z., G.W., G.S., K.T., M.B., E.T. designed and performed the experiments in this study and analyzed the data. M.V. and J.S. helped with experiments related to SF3B1. H.C. helped with the experiment design for MEK1 studies. B.E.B. provided additional supervision. X.D.C., Z.C., B.E.B., and F.C. wrote the manuscript with help from all authors.

## Data and materials availability

Sequencing data will be available at Sequence Read Archive. Expression plasmids will be made available on Addgene under UBMTA. The code for this paper is available at https://github.com/chen-dawn/hace.

