## Supplementary Information for "Helicase-assisted continuous editing for programmable mutagenesis of endogenous genomes"

**This PDF file includes:**

Materials and Methods
Supplementary Figure 1 to 7
Supplementary Tables 1 to 13
References

### Materials and methods

Plasmids and oligonucleotides

The guide sequences used for HACE mutagenesis (Supplementary Table 1) are closed by Gibson assembly or Golden Gate assembly. The oligos used in this study for sequencing (Supplementary Table 2) were purchased from Integrated DNA technologies (IDT) or Azenta/GENEWIZ. The Cas9 nickase plasmids were derived from plasmids pSpCas9(BB)-2A-GFP (Addgene 48138) and pCMV-PEmax-P2A-GFP (Addgene 180020). Plasmids expressing sgRNAs and pegRNAs were cloned by Gibson assembly or Golden Gate assembly. HACE Editor plasmids were cloned by Gibson assembly of PCR products. Individual helicases were either subcloned from plasmids (pEGFP-BLM - Addgene 110299; pET22B_SA_PcrA - Addgene 102999; pCMV-Tag1-NS3 - Addgene 17645) or synthesized by Integrated DNA Technologies after mammalian codon optimization. The helicases tested are summarized in Supplementary Table 3, and sequences of individual helicases tested are listed in Supplementary Table 4. All new plasmids generated during this study will be deposited on Addgene.

Mammalian cell culture

HEK293FT cells (Thermo Fisher - R70007) and A375 cells (ATCC, CRL-1619) were cultured in Dulbecco's Modified Eagle Medium with GlutaMAX (Thermo Fisher Scientific 10564011) supplemented with 10% (v/v) fetal bovine serum (FBS, Sigma-Aldrich F4135) and 1x penicillin-streptomycin (Thermo Fisher Scientific 15140122). Adherent cells were maintained at confluency below 80%-90% at 37℃ and 5% $CO_2$.

K562 cells (ATCC, CCL-243) were cultured in RPMI 1640 medium with GlutaMax (Thermo Fisher-61870036) supplemented with 10% (v/v) FBS and 1x penicillin-streptomycin. Suspended cells were maintained at confluency below $1.5x10^6$ cells/ml at 37℃ and 5% $CO_2$. For stimulation experiments, 50ng/ml Phorbol12-myristate13-acetate (PMA, Sigma-Alrich, P8139) and 500ng/ml ionomycin calcium salt from Streptomyces Conglobatus (ionomycin, Sigma-Aldrich, I0634) are added to the cell culture media to simulate the cells for 2-3 hours.

Transfection of HACE plasmid and genomic DNA preparation for HACE editing assays

The day before transfection, 10,000 HEK293FT cells were seeded per well on 96-well plates (Corning). Then, 16-24 hours after seeding, cells were transfected at approximately 70% confluency with 0.3uL of TransIT-LT1 (Mirus Bio) according to the manufacturer's specifications. Each well was transfected with 40ng of HACE editor plasmid, 40ng of Cas9 nickase plasmid, and 16ng of sgRNA plasmid were delivered to each well unless otherwise specified. For control conditions, HACE editor plasmid and/or Cas9 nickase plasmid were substituted with the same amount of pUC19 plasmid. Cells were cultured for 3 days after transfection. DNA was collected from transfected cells by removal of medium, resuspension in 50 uL of QuickExtract (Lucigen), and incubation at 65ºC for 15 min, 68ºC for 15 min, and 98ºC for 10 min. After thermocycling, lysate was used directly in downstream PCR reactions as per manufacturer protocol.

High-throughput DNA sequencing of genomic DNA samples

For short amplicon (~300bp) sequencing for HEK293T and A375 cells, the target region was amplified from genomic DNA samples using Phusion U Hot Start PCR master mix (Thermo

Fisher Scientific, F562) in a 20uL reaction. The following program was used: 98ºC for 30 s; 28 cycles of 98ºC for 10 s, 65ºC for 30 s, 72ºC for 30 s; 72ºC for 2 min, then 4ºC forever. Barcodes and adapters for Illumina sequencing were added in a subsequent PCR amplification using Q5 High-Fidelity Hot-Start Polymerase Master Mix (2x, New England Biolabs). Amplicons were pooled and prepared for sequencing on a NextSeq (Illumina) with paired-end reads (read1,160bp; index1, 8bp; index2, 8 bp; read2, 160bp). Reads were demultiplexed and analyzed with appropriate pipelines.

For long amplicon (~2000 bp) sequencing, the targeted region was amplified using Phusion U Hot Start PCR master mix in a 20uL reaction. The following program was used: 98ºC for 30 s; 28 cycles of 98ºC for 10 s, 65ºC for 30 s, 72ºC for 2 min; 72ºC for 5 min, then 4ºC forever. PCR products were purified using Magnetic Ampure XP beads (Beckman Coulter) using a 1:1 bead solution:DNA solution ratio to select the PCR fragments. Purified PCR products were eluted in 20uL of water. The concentration of each sample was measured by Qubit (Thermo Fisher Scientific). The sequencing library was prepared following the Nextera XT Kit protocol (Illumina) using 1ng of purified amplicon DNA per sample as starting material and half of the recommended amount of each kit reagent. Sequencing was performed on a NextSeq (Illumina) with paired-end reads (read1,100bp; index1, 8bp; index2, 8bp; read2, 100bp).

Quantification of editing rate

Raw fastq reads obtained from sequencing were quality trimmed using BBduk[1] (BBMap v38.93) with the options "qtrim=rl trimq=28 maq=25". Next, all bases with quality scores below 28 were masked to N using seqtk v1.3[2]. The filtered reads were aligned to the reference sequence using Bowtie2[3] (version 2.3.4.3). The pileup at each base was calculated using a custom Python script.

To calculate the mutation rate, we filtered for base positions with a sequencing coverage of at least 10,000. Bases that had a higher than 5% mutation rate in the control condition were masked since this either indicated that it was a variant or an artifact from sequence alignment. The average G>A editing rate was calculated by extracting all positions where "G" was the reference base, then taking the average of the per base G>A editing rate. The editing rate for other base transition and transversion modes was calculated similarly.

To calculate the local editing rate, the alignment was centered such that the nick site is centered at base position 0. For every base position, the local G>A editing rate was calculated by extracting all the "G" bases within a 100bp window (50bp upstream and 50bp downstream) and then taking the average of all per G>A editing rates. Code for quantifying the edit rate is available at https://github.com/chen-dawn/hace.

Cell viability measurements

Cell viability was measured using the CellTiter-Glo Luminescent Cell Viability Assay (Promega) following the manufacturer's protocols. Briefly, HEK293FT cells were seeded at a density of 10,000 cells per 100uL per well in a 96-well plate in biological triplicates. The following day, cells were transfected with respective HACE plasmids according to the above protocol. Cell viability was measured 72 h after transfection. Luminescence readings were performed using a SpectraMax M5 (Molecular Devices) plate reader.

Whole exome sequencing and off-target analysis

The day prior to transfection, 50,000 HEK293FT cells were seeded in a 24-well plate. The following day, individual HE constructs were transfected together with a sgRNA targeting the *MAP2K1* locus. Genomic DNA was extracted from cells 3 days post-transfection using the Zymo Quick-DNA Miniprep Kit (Cat D3024). Amplicon sequencing was performed at the *MAP2K1* loci to confirm that there is HACE-dependent editing at the target loci in each condition. The whole genome DNA sequencing library was prepared using the NEBNext Ultra II FS DNA Library Prep Kit for Illumina (Cat E7805S). Exome sequences were enriched using the xGen Exome Hybridization Panel (IDT 10005152) following the manufacturer's protocols. Exome libraries were sequenced on a NovaSeq X (Illumina) with paired-end reads (read1,150bp; index1, 8bp; index2, 8 bp; read2, 150bp) at a minimum of 100 million reads per sample.

The sequencing output was demultiplexed using bcl2fastq, and the paired-end reads were aligned to the reference genome hg38 using HISAT2 v2.2.1[4]. Aligned reads from each replicate were subsampled using reformat.sh (BBMap v38.93) and 100 million aggregated reads per replicate for each condition were used for further analysis. The HEK293FT-specific single-nucleotide polymorphisms (SNPs) were determined following the GATK4 variant calling workflow for germline short variant discovery (https://gatk.broadinstitute.org/hc/en-us/articles/360035535932-Germline-short-variant-discovery-SNPs-Indels-) on wild-type HEK293FT exome libraries (>50x coverage). In brief, the aligned reads were de-duplicated using Picard v2.27.5. HaplotypeCaller (GATK4) was used for calling variants, and known variants in dbSNP version 138 were used during base-quality recalibration. The chromosomal coordinates where SNPs were detected were excluded from subsequent analysis.

To quantify the per-base editing rate of the exome, the base pileup at each base was calculated using samtools mpileup (v 1.15.1), followed by post processing using mpileup2readcounts (https://github.com/IARCbioinfo/mpileup2readcounts). Bases with less than 50 total read depth were excluded from subsequent analysis. The genome was binned into 100kb bins using bcftools v1.15.1. The off-target C>T editing rates for each genomic bin were obtained using a custom R script by counting the number of C and T bases in each bin. Fisher's exact test was used to quantify significant changes in editing for each bin relative to cells transfected with only nCas9, using the FDR correction to adjust for multiple hypothesis testing. Significant off-target sites are listed in Supplementary Table 5.

MEK1 inhibitor-resistance screen

A375 cells were diversified for 3 days by transfection of HE variant AID-PcrA M6-UGI, nCas9 D10A, and sgRNAs targeting exons 2, 3, and 6 of the *MEK1* gene using TransIT-2020 (Mirus Bio). Approximately 5 million cells in a 15-cm dish were placed under selection with either 100nM selumetinib or 5nM trametinib for 20 days. A portion of pre-selection cells were harvested as a control. Cells were passaged every 3 days to ensure they were maintained at <70% confluency. After selection, cells were harvested, and genomic DNA was extracted using QuickExtract (Lucigen). The *MEK1* exons were amplified with exon-specific primers (Supplementary Table 2) using Phusion U Hot Start Master Mix. Concurrently, we also harvested RNA from selected cells using the Qiagen RNeasy Mini Plus Kit (Cat 74134). The cDNA was generated by reverse transcription using Maxima H Minus Reverse Transcriptase (Thermo

Fisher). Sequencing libraries for cDNA were generated using the modified Nextera XT Kit protocol described in the "High-throughput DNA sequencing of genomic DNA samples" section above. All libraries were sequenced on a NextSeq (Illumina) with paired-end reads (read1, 160 bp; index1, 8 bp; index2, 8 bp; read2, 160 bp). The mutation rate (allele frequency) for each base of the *MEK1* sequence was calculated for both pre-and post-selection samples. Significant mutations were identified by comparing the base counts between pre- and post-selection samples using a Fisher's exact test (Supplementary Table 6). We compared the mutation rate between RNA and DNA samples and found that they had a high correlation.

SRE reporter assay

pEF1a-*MEK1* wild type, pEF1a-*MEK1*G128D, pEF1a-*MEK1*G202E, and pEF1a-*MEK1*E203K were generated using Gibson assembly. Sequences of *MEK1*-derived constructs are available in Supplementary Table 7. The SRE reporter assay was performed using the SRE reporter kit (BPS Biosciences) according to the manufacturer's protocols. In brief, ~10,000 HEK293FT cells in 100 µl of growth medium were seeded in 96-well white opaque assay plates. The cells were transfected with 60 ng of reporter plasmid and 40 ng of respective *MEK1* plasmids. The culture medium was replaced 6 hours post-transfection with 50 µl of trametinib-containing medium with 0.5% FBS. After 12 hours, the cells were washed and incubated with 50 µl of 0.5% FBS-containing culture medium supplemented with recombinant human epidermal growth factor protein (Life Technologies) at a final concentration of 10 ng/mL. After 6 hours of incubation, the reporter activity was assayed using a dual luciferase (Firefly-Renilla) assay system (BPS Bioscience) according to the manufacturer's instructions using a SpectraMax M5 (Molecular Devices) plate reader. The ratio between Firefly luminescence and Renilla luminescence intensity was calculated for each well after background subtraction.

Design of *SF3B1* splicing minigene reporter

The minigene reporter to probe SF3B1 function was constructed by Gibson assembly of a synthetic minigene sequence (synthesized by Twist Biosciences) into a custom bicistronic mCherry/GFP reporter plasmid. To construct the minigene reporter, we fused the VCP exon 10 sequence and 150 bp of its immediate downstream intron with DLST exon 6 and 97 bp of its immediate upstream intron. We also appended an "ATG" start codon at the beginning of the sequence. The open reading frame was adjusted such that correct splicing in wild-type cells will result in pre-mature termination before the GFP. In contrast, the alternative 3' splice-site usage in *SF3B1* mutant cells will result in full-length GFP expression. The minigene reporter sequences are annotated in Supplementary Table 8.

*SF3B1* mis-splicing screen

HEK293FT cells were diversified for 3 days by co-transfection of HE variants AID-PcrA M6-UGI and TadA-PcrA M6-UGI, nCas9 D10A, sgRNAs targeting *SF3B1* exons 13-17, and splicing mingene reporter. Cells transfected with only minigene reporter were used as undiversified control. The experiment was performed in triplicate, with ~10 million cells transfected per replicate. After diversification, cells were prepared for flow sorting by washing and resuspending in 1x PBS with 2% BSA. Cells were sorted using a SONY MA900 sorter, where mCherry-positive cells were sorted into a $GFP^-$ and $GFP^+$ bin. At least 1 million cells were collected for each cell population. After flow sorting, the RNA of the cells was extracted using the Qiagen RNeasy Mini Plus Kit. The cDNA was generated by reverse transcription using

Maxima H Minus Reverse Transcriptase. Sequencing libraries for cDNA were generated using the modified Nextera XT Kit protocol and sequenced on a NextSeq. Fold enrichment was calculated by dividing the mutation rate in $GFP^+$ by that of $GFP^-$ samples. The significant mutations were identified using a Fisher's exact test and are shown in Supplementary Table 9. The clinical mutations that are observed in *SF3B1* were retrieved from COSMIC[5]. A mutation was considered high frequency if there were at least 3 observations in the dataset.

*CD69* enhancer tiling for functional bases

K562 cells were nucleofected with 2.5 µg of HE and 2.5 µg of nCas9 and sgRNA plasmids using the SF Cell Line 4D-Nucleofector X Kit L (Lonza V4XC-2024), following the manufacturer's protocol. Each plasmid contained a fluorescent protein reporter (sgRNA mCherry, nCas9 GFP, HE BFP). Approximately $1.5\text{-}2\times10^6$ cells were used per nucleofection reaction. After 24 hours, cells were sorted using either SONY SH800 or BD Aria flow cytometry sorter to isolate cells expressing all plasmid components.

On day 7 post-nucleofection, cells were stimulated with PMA/Ionomycin for 2-3 hours. The cells were stained with the antibody cocktail in the staining buffer of a 1:1 mix of PBS and Brilliant Staining Buffer (BD 566349) at room temperature for 20 mins or at 4°C for 30 mins. The following antibodies and dyes from BioLegend were used: Brilliant Violet 510 anti-human CD69 Antibody (310936); APC anti-human CD69 Antibody (310910); Zombie NIR Fixable Viability Kit (423106). Cells were washed once in PBS with 1%FBS and then resuspended in the same buffer to prepare for flow sorting. Subsequently, the top 40% of cells showing high CD69 expression ($CD69^{high}$) and the bottom 20% with low CD69 expression ($CD69^{low}$) were sorted using the SONY SH800 flow cytometer. A minimum of 100,000 cells were collected per tube. Genomic DNA was then isolated from these cells either by using the QIAGEN DNA micro isolation kit (Cat #56304) or by lysis buffer (0.5% Triton X-100, 0.1 AU/ml QIAGEN Protease (Cat 19157) in $H_2O$). The lysis process involved incubation at 56°C for 20 minutes and at 72°C for 20 minutes at 600 rpm on a thermo shaker. Amplicon PCR for the genomic DNA was processed using the KAPA HiFi HotStart ReadyMix PCR Kit (Roche, KR0370). The following program was used: 95℃ for 5 min; 30 cycles of 95℃ for 30 s, 60℃ for 30 s, 72℃ for 30 s; 72℃ for 5 min; 4℃ forever. The amplicon libraries were sequenced on a NextSeq.

To identify enriched bases, the %C->T or %G->A of each group were first calculated for both $CD69^{high}$ and $CD69^{low}$groups (%high or %low). Then the log2 odds ratio of $CD69^{high}$ versus $CD69^{low}$ was calculated as $\log_2 OR = \log2\ [(\%high/(1\text{-}\%high))/(\%low/(1\text{-}\%low))]$. The correlation of technical replicates was plotted using GraphPad Prism 10.0. The top hits are recorded in Supplementary Table 10.

Base editing validation

The following base editors are used in this study: pRDA_478 (Addgene 179096), pRDA_479 (Addgene 179099), pCAG-CBE4max-SpG-P2A-EGFP (Addgene: RTW4552/139998), pCAG-CBE4max-SpRY-P2A-EGFP (Addgene: RTW5133/139999), pCMV-T7-ABE8.20m-nSpCas9-NG-P2A-EGFP (Addgene: KAC1164/185919), pCMV-T7-ABE8.20m-nSpRY-P2A-EGFP (Addgene: KAC1335/185917). The validation sgRNAs are listed in Supplementary Table 11. The sgRNA sequences were cloned into pCMV-BFP-U6-sgRNA (Addgene:196725, gift from Bernhard Schmierer) or directly into pRDA_478 or pRDA_479. Mutations for sgRNAs and bystander editing rates were quantified using CRISPResso2[6].

To validate *MEK1* variants, 67 ng of cytidine base editor and 33 ng sgRNA plasmids were transfected into A375 cells per well in a 96-well format. A non-targeting sgRNA was used as a control. Post-diversification for 3 days, the cells were selected with either 100 nM selumetinib or 5 nM trametinib for 14 days. The mutation rate pre- and post-selection were analyzed by amplicon sequencing. All experiments were conducted in triplicates.

For validation of *SF3B1* variants, HEK293FT cells in a 96-well format were transfected with 67 ng of the base editor-sgRNA plasmid and 33 ng of the minigene splicing reporter per well. A non-targeting sgRNA was used as a control. Cells were diversified for 3 days, then the GFP:mCherry ratio in each well was quantified by confocal microscopy using a custom cell segmentation and quantification pipeline. Briefly, individual cells were segmented via watershed segmentation using the mCherry channel. For each segmented cell, the total pixel area and mean intensity of the pixels were computed for GFP (488 nm) and mCherry (561 nm) channels to obtain a "pseudo-flow cytometry" dataset. The fluorescence background for each channel was subtracted from all conditions in that channel, and aggregated values for each condition were divided by area to obtain average fluorescence intensity. Standard deviation was computed by comparing average values in three technical transfection replicates. Editing at each sgRNA was quantified by amplicon sequencing of genomic DNA samples from each well. All experiments were conducted in triplicates.

For validation of *CD69* variants, 2 µg of the base editor plasmid and 2 µg of the sgRNA plasmid were nucleofected into 1.5 x $10^6$ K562 cells using the SF Cell Line 4D-Nucleofector X Kit L according to the manufacturer's protocol. At 24 h post nucleofection, the cells co-expressing base editor and sgRNA were sorted based on reporter expression. On day 4 post-nucleofection, cells were stimulated with PMA/ionomycin for 2-3 hours, and the top 40% of CD69 high expression cells and bottom 20% of CD69 low expression cells were sorted using a Sony SH800 flow cytometer, collecting at least 10,000 cells per tube. Genomic DNA was isolated from the sorted cells using the QIAgen DNA Micro Kit (Cat# 56304) and prepared for amplicon sequencing using the protocol described in the "High-throughput DNA sequencing of genomic DNA samples" section. The mutation rate at each locus was quantified using CRISPResso2.

<u>Prime editing validation</u>
The following prime editor plasmids are used in this study:pCMV-PEmax-P2A-hMLH1dn (Addgene: 174828), pCMV-PEmax-P2A-GFP(Addgene:180020), pEF1a-hMLH1dn (Addgene:174824). Desired pegRNA and nickase sgRNA sequences were designed using PrimeDesign[7]. The epegRNA overhang was designed using pegLIT[8]. Sequences of pegRNAs are shown in Supplementary Table 12.

For prime editing validation of *SF3B1* variants, HEK293FT cells in a 96-well format were transfected with 150 ng of PEmax, 50 ng of epegRNA, 25 ng of nicking sgRNA, and 50 ng of minigene splicing reporter using 0.5uL of TransIT-LT1 per well. Cells were diversified for 3 days, then the GFP:mCherry ratio in each well was quantified by confocal microscopy as described above.Editing at each sgRNA was quantified by amplicon sequencing of genomic DNA samples from each well using CRISPResso2.

For prime editing validations for *CD69* enhancer variants in K562 cells, 2ug of the prime editor plasmid, 1ug of hMLH1dn plasmid and 1ug of epegRNA plasmid, and 0.5ug of nickase sgRNA plasmid were nucleofected in 1.5 x $10^6$ cells using SF Cell Line 4D-Nucleofector X Kit L according to the manufacturer's protocols. After 24 hours, the cells that were positive for both prime editor and epegRNA were sorted based on the GFP and mCherry reporters and cultured in regular complete RPMI media. A second round of nucleofection and sorting was performed 4 days post-transfection to increase prime editing efficiency. On day 5 post the second nucleofection, CD69 expression levels were quantified by flow cytometry. Genomic DNA was harvested from $CD69^{high}$ (top 40%) and $CD69^{low}$ (bottom 20%) cells, and the editing efficiency was quantified by performing amplicon sequencing using the protocols described above. The mutation rate at each locus was quantified using CRISPResso2.

**Supplementary Figure 1.**

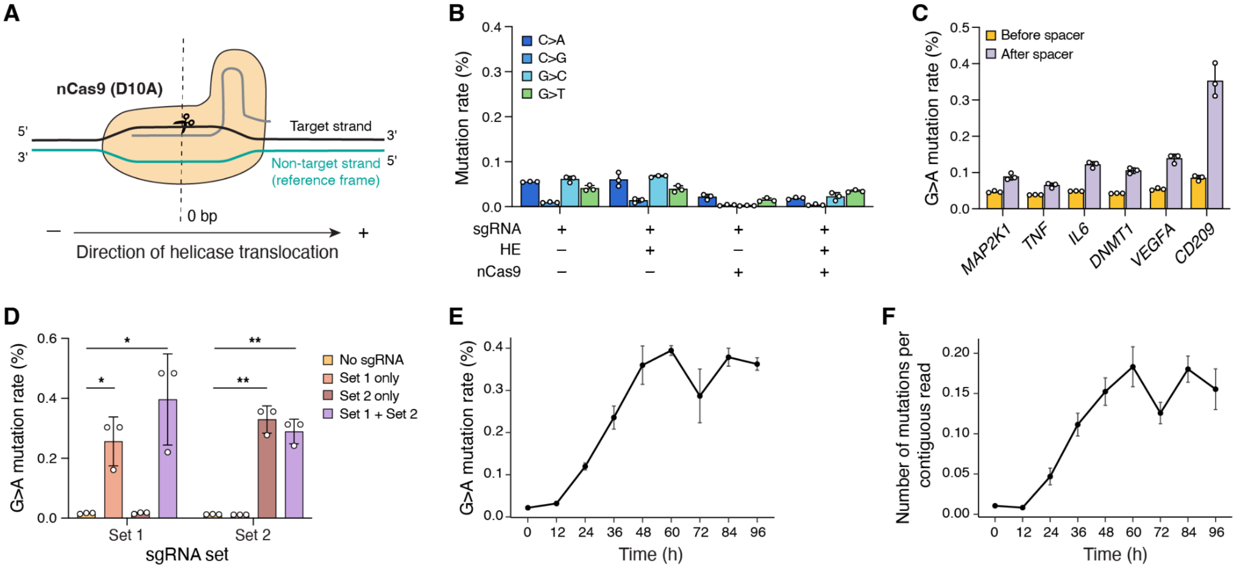

**Supplementary Figure 1 Characterization of the HACE system**
**(A)** Schematic of  direction of helicase translocation relative to position of sgRNA. Vertical dashed line represents the location of the nick. The non-target strand (DNA strand that does not bind sgRNA) is depicted in green. The helicase translocates in the 3’ to 5’ direction relative to the non-target strand. **(B)** Average mutation rate across diverse base transition and transversion modes for the region downstream of the nick site. **(C)** Average G>A mutation rates before and after sgRNA spacer. **(D)** G>A mutation rate for two sets of three sgRNAs each. Significance is determined via paired two-tailed *t*-test between the presence and absence of sgRNAs. $*P < 0.05$. $**P < 0.01$. **(E)** G>A mutation rate over the course of 96 h with transfected HE, nCas9, and sgRNA. **(F)** Mean mutations per contiguous read over time points. All data are mean of technical replicates (n=3) ± s.d.

**Supplementary Figure 2.**

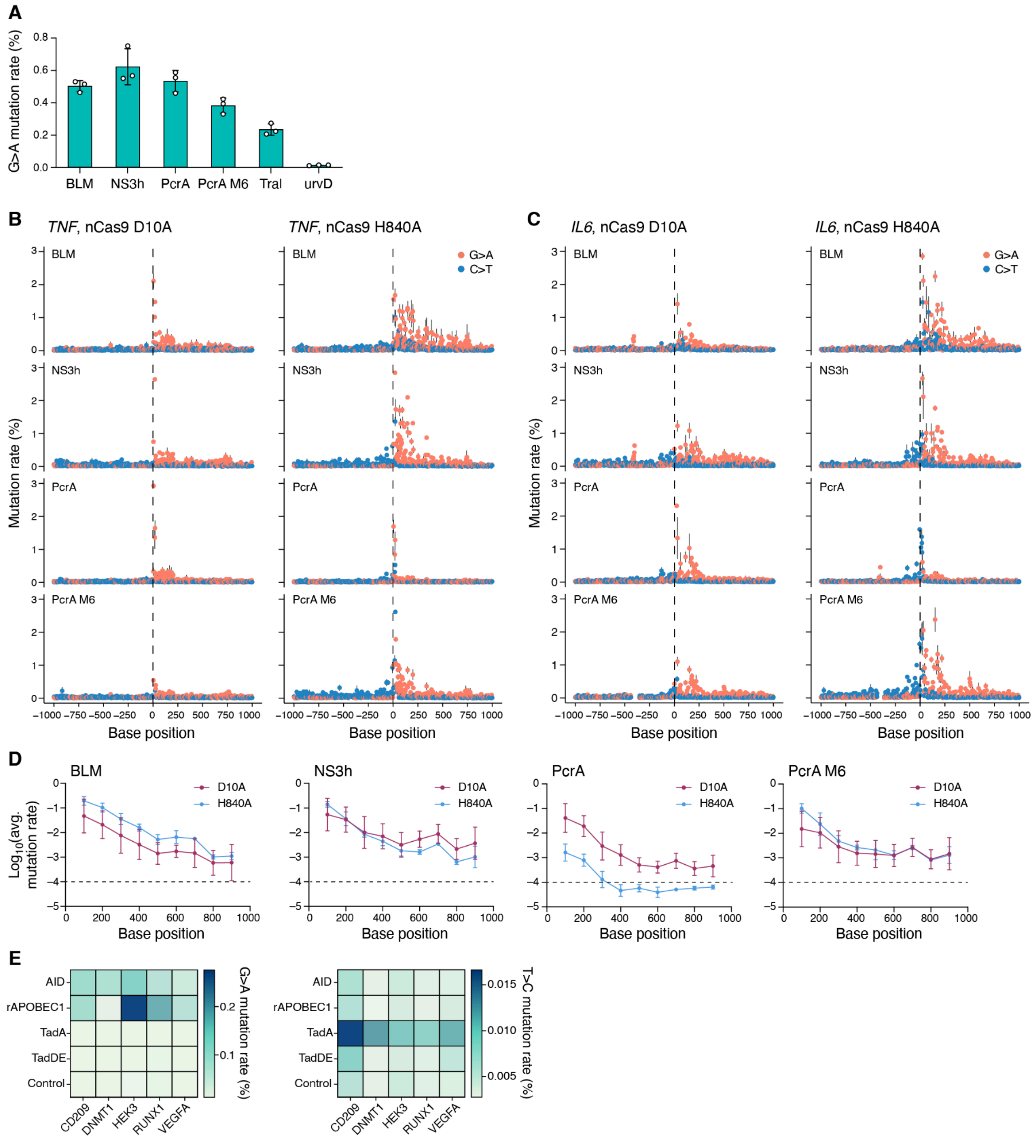

**Supplementary Figure 2 HACE has long range and activity across diverse helicases and deaminases.**

**(A)** Mutation rate at the *HEK3* loci for HEs with different helicase variants and the nCas9(D10A) nickase variant. Data are mean of technical replicates (n= 3) ± s.d. **(B)** Mutation rate per base across a ~1-kb target region at the *TNF* loci for HEs with different helicase variants and either the nCas9(D10A) or nCas9(H840A) nickase variant. The vertical dashed line shows the nick site.

Data are mean of technical replicates (n= 3) ± s.d. **(C)** Mutation rate per base across a ~1-kb target region at the *IL6* loci for HEs with different helicase variants and either the nCas9(D10A) or nCas9(H840A) nickase variant. The vertical dashed line shows the nick site. Data are mean of technical replicates (n= 3) ± s.d. **(D)** The local mutation rate per every 100 bp window across a ~1-kb region from the nick site for HEs with different helicase variants and either the nCas9(D10A) or nCas9(H840A) nickase variants. The local mutation rate is the average across 3 target loci. **(E)** Average G>A and T>C mutation rate at 5 different genomic loci for HEs with different deaminase variants fused to the PcrA M6 helicase.

**Supplementary Figure 3.**

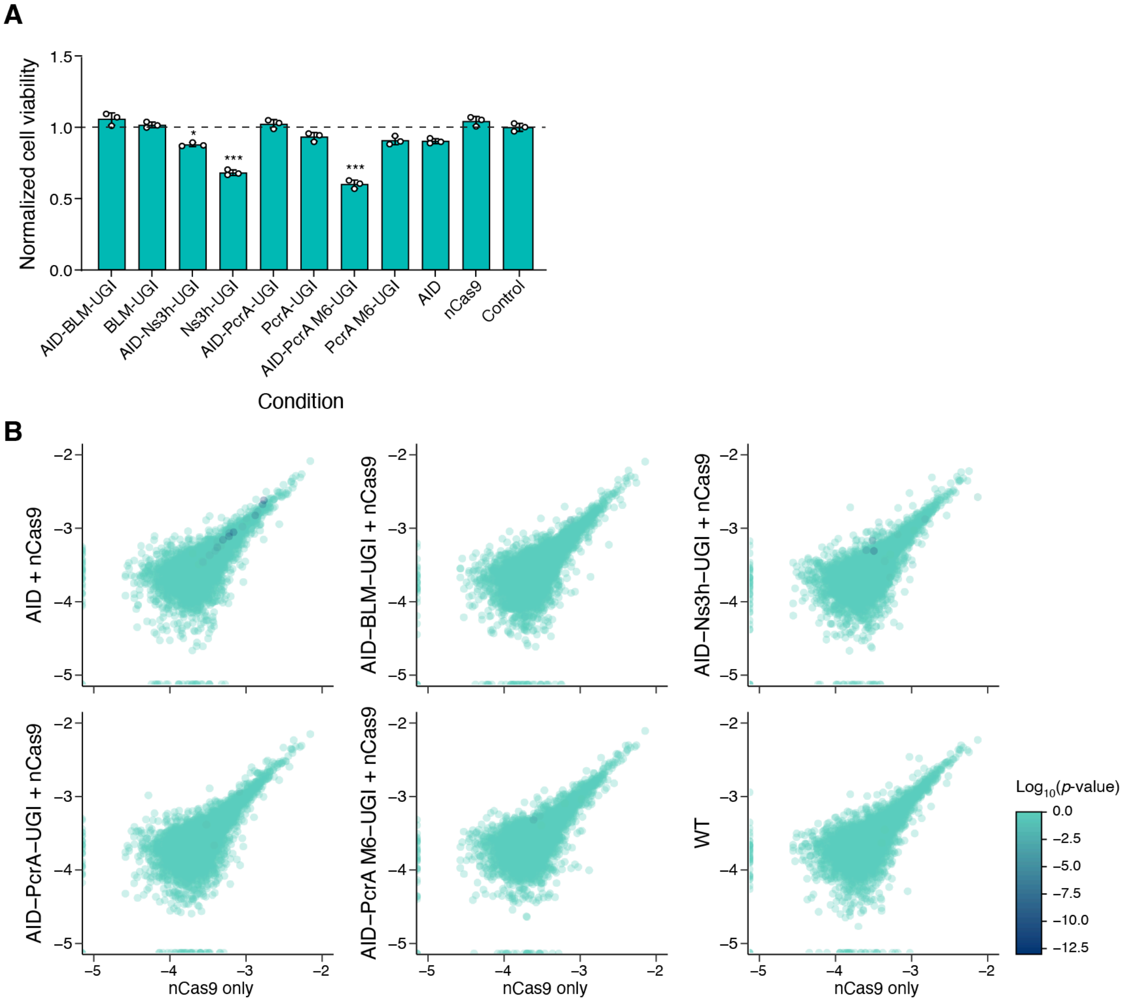

**Supplementary Figure 3 Evaluation of HACE toxicity and off target editing**
**(A)** Cell viability of HEK293FT cells after transfection of HEs with different helicase variants or AID alone. All experiments were transfected with nCas9(D10A) and a sgRNA targeting the *MAP2K1* loci. Data are mean of technical replicates (n= 3) ± s.d. P-values for conditions with significant difference between control and treated groups (unpaired two-tailed *t*-test) are annotated above the bars. $*P < 0.05$. $***P < 0.001$. **(B)** Analysis of exome-wide off-target editing. Scatter plots show the average C>T mutation rate for 100kb genomic bins in cells transfected with AID alone or HE constructs with different helicases compared with control cells transfected with nCas9(D10A) only. Sites are colored by FDR-adjusted *P* value (color bar, left). Experiments were generated from two independent replicates.

**Supplementary Figure 4.**

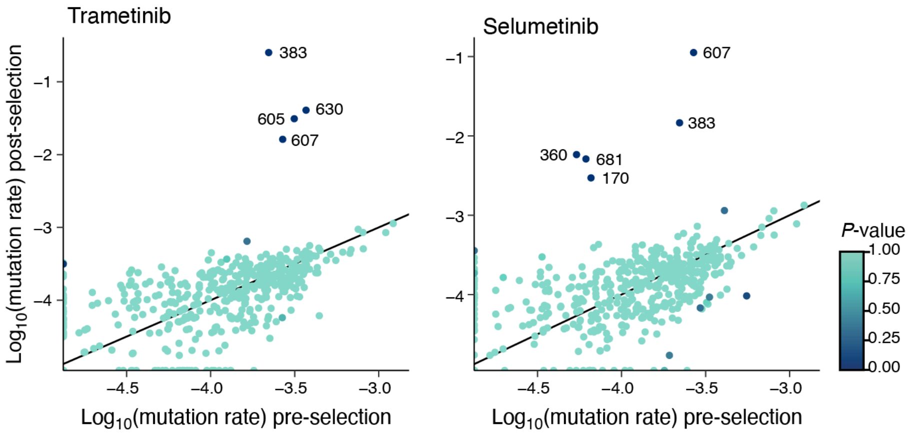

**Supplementary Figure 4 Identification and validation of mutations leading to MEK1-inhibitor resistance.**

Scatter plot shows the mutant vs. reference allele frequency for A375 cells selected with either trametinib (left) or selumetinib (left) compared to control cells. Sites are colored by Bonferroni-corrected *P* value (color bar, left). Significant CDS base positions are annotated.

**Supplementary Figure 5.**

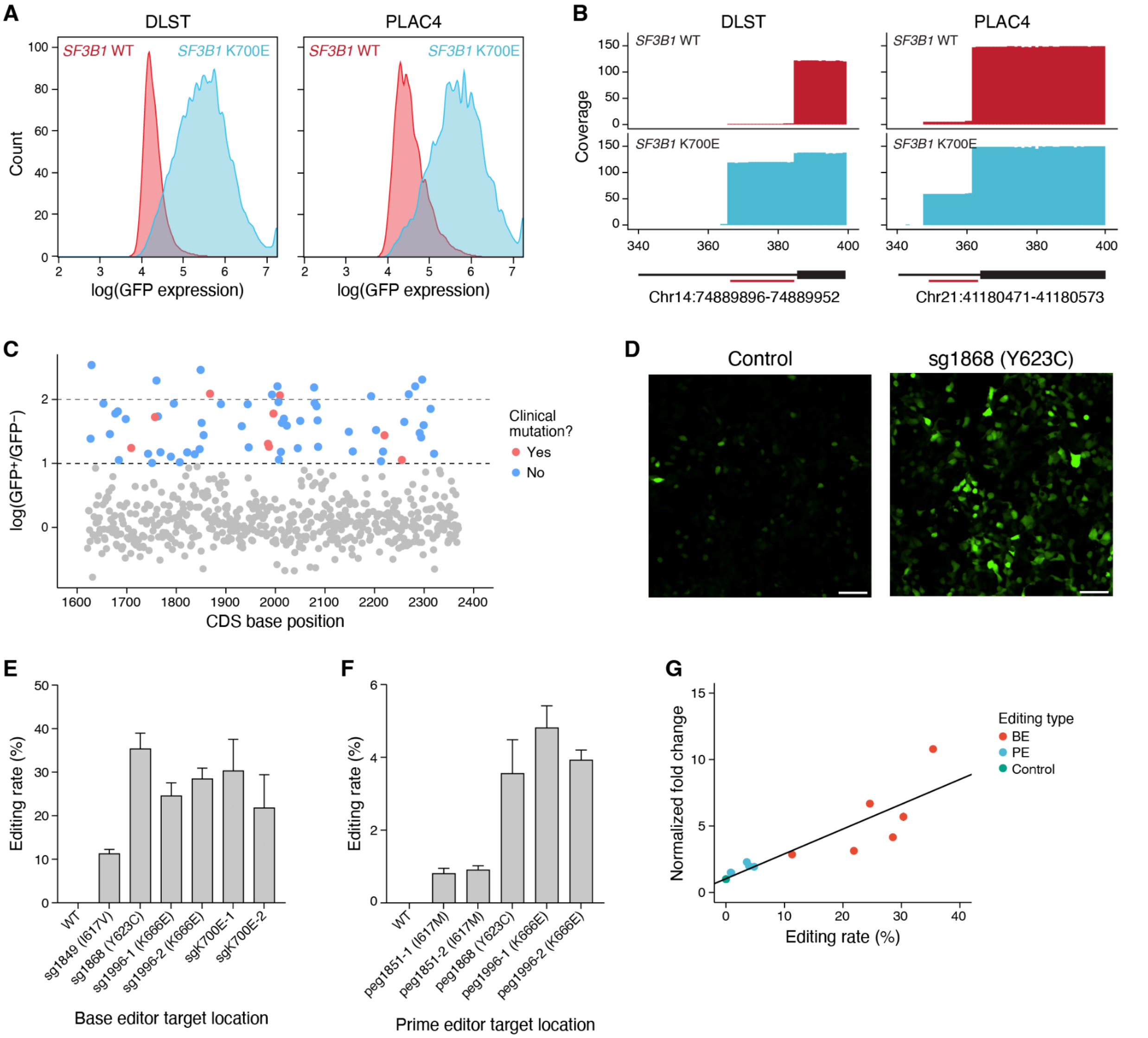

**Supplementary Figure 5 SF3B1 minigene reporter and validation of SF3B1 mutations via base and prime editing.**

**(A)** Histogram of GFP signal measured by flow cytometry between isogenic K562 $SF3B1^{WT}$ and $SF3B1^{K700E}$ cells for two different minigene reporter constructs. Cells were gated for mCherry expression. **(B)** RNA base pileup for 2 minigene reporter constructs nucleofected into isogenic K562 $SF3B1^{WT}$ and $SF3B1^{K700E}$ cells. The location of intron-exon junctions is annotated in black. The sequence that is retained by alternative 3′ss is annotated in red. **(C)** Fold enrichment of SF3B1 cDNA sequence in $GFP^{+}$ vs $GFP^{-}$ samples. The mutations with >10-fold base enrichment are colored by whether they have been observed >=3 times in clinical samples. **(D)** representative images of minigene GFP reporter expression in control cells vs cells with candidate mutations introduced by base editing (sg1668, Y623C). **(E)** Editing rate in mutations installed via base editing. Data are mean of technical replicates (n= 3) ± s.d. **(F)** Editing rate in mutations installed via prime editing. Data are mean of technical replicates (n= 3) ± s.d. **(G)**

correlation of editing rate to normalized fold-change for minigene reporter ($\rho = 0.880$). Data are mean of technical replicates ($n = 3$) ± s.d.

**Supplementary Figure 6.**

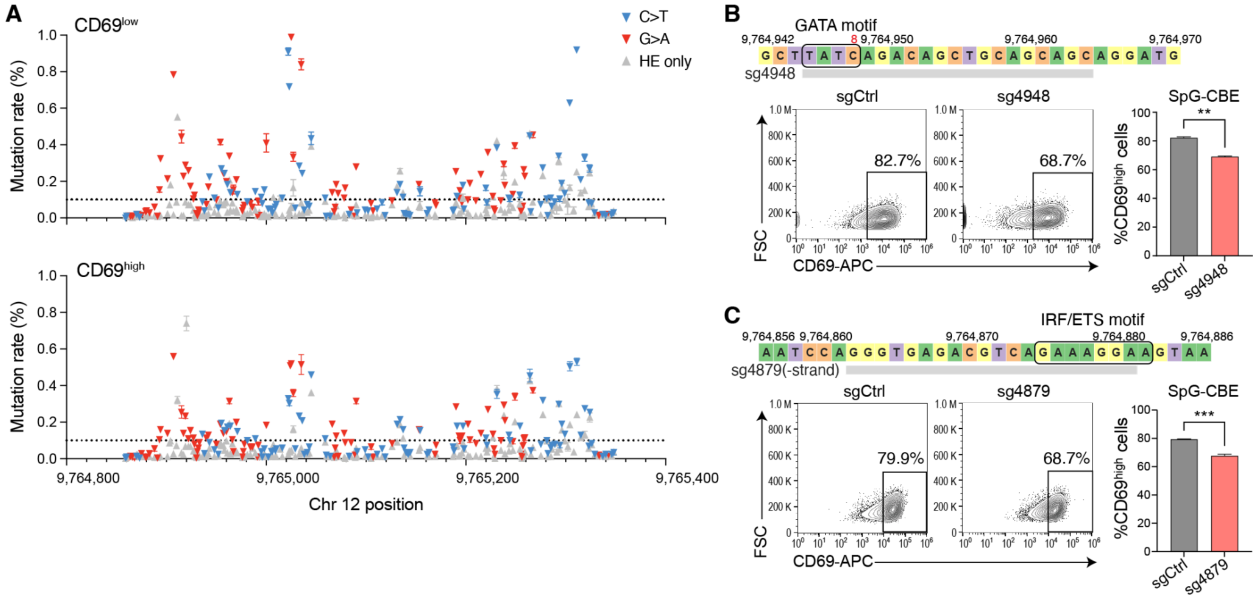

**Supplementary Figure 6 HACE mutagenesis and validation on the CD69 enhancer region.** **(A)** Per base mutation rate across the core region of CD69 enhancer for CD69$^{low}$ and CD69$^{high}$ sorted populations. G>A and C>T transitions are colored red and blue, respectively. Grey dots represent the editing rate from control groups. Each data point represents mean ± s.e.m. (n=3). **(B)** Sequence of the sg4948 target site, with the GATA motif boxed (top). Proportion of CD69$^{high}$ cells after stimulation on day 7 post-transfection in base-edited cells using sg4948 compared to control cells as quantified by flow cytometry (bottom). **(C)** Sequence of the sg4879 target site, with the GATA motif boxed (top). Proportion of CD69$^{high}$ cells after stimulation on day 7 post-transfection in base-edited cells using sg4879 compared to control cells as quantified by flow cytometry (bottom). Significance is determined via unpaired two-tailed *t*-test between control and edited groups. ***P* <0.01. ****P* < 0.001.

**Supplementary Figure 7.**

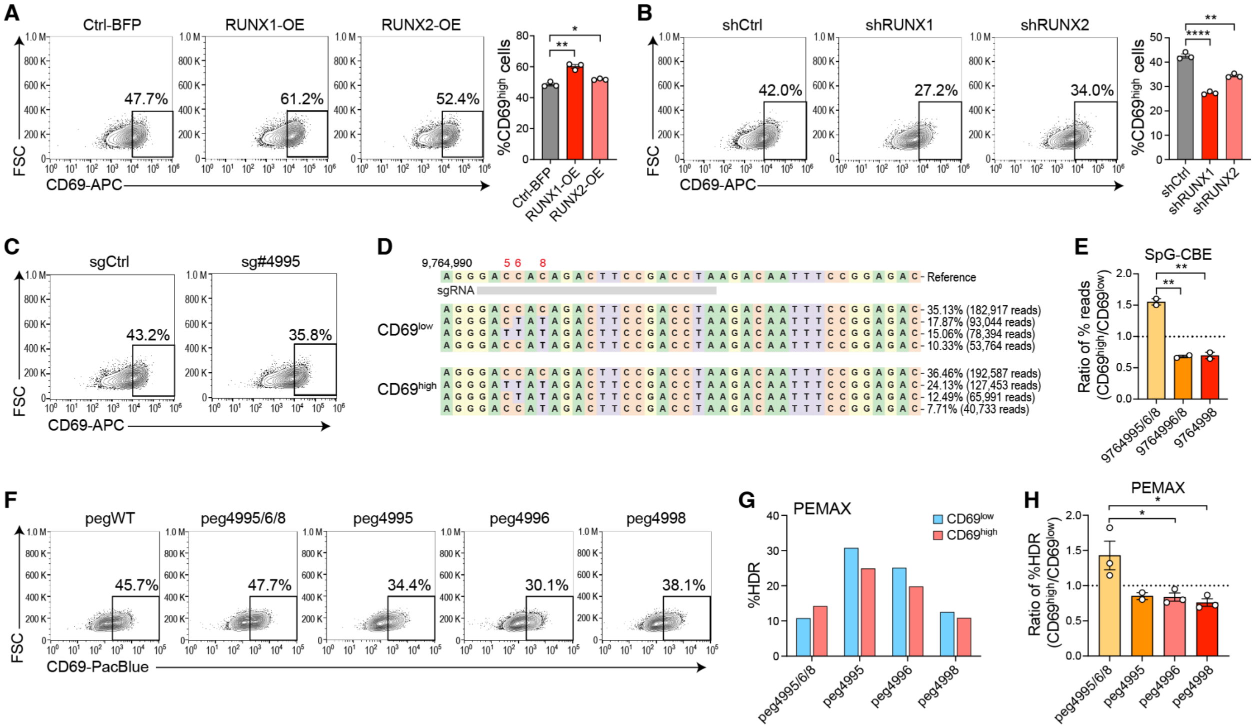

**Supplementary Figure 7 RUNX1/2 regulates CD69 expression via the CD69 enhancer.** **(A)** Flow cytometry and bar plot depict the proportion of $CD69^{high}$ cells after stimulation in control, RUNX1-overexpression (OE), and RUNX2-OE groups three days post-transfection. Data are from 2 independent experiments, each with 3 technical replicates, mean ± SEM. **(B)** Flow cytometry and bar plot depict the proportion of $CD69^{high}$ cells after stimulation in control, RUNX1-shRNA, and RUNX2-shRNA groups three days post-transfection. Data are from 2 independent experiments, each with 3 technical replicates, mean ± s.e.m.. **(C)** Proportion of $CD69^{high}$ post-stimulation for cells targeted with control sgRNA (sgCtrl) or sg4995 with SpG-CBE base editor 4 days post-transfection. **(D)** Top alleles with different combinations of C>T mutations in cells targeted with base editing and sg4995 as quantified by amplicon sequencing in both $CD69^{low}$ and $CD69^{high}$ populations. **(E)** Ratio of $CD69^{high}$ to $CD69^{low}$ for sequencing read proportions for different base edit combinations as depicted in (D). **(F)** Representative flow cytometry plots depict the proportion of $CD69^{high}$ after stimulation in cell populations targeted with different epegRNAs using prime editing. **(G)** Frequency of perfect homologous recombination rate (%HDR) in cell populations targeted with different epegRNAs using prime editing. **(H)** Ratio of %HDR between $CD69^{high}$ and $CD69^{low}$ for editing quantified in (G). For all comparisons, significance is determined via an unpaired two-tailed *t*-test between control and edited groups. *$P$ <0.05. **$P$ <0.01. ***$P$ < 0.001.

**Supplementary Table 1. Table of guide sequences for HACE.**

| sgRNA name | Target gene/region | Experiment | Spacer sequence |
|---|---|---|---|
| HEK3.1 | HEK3 | Initial validation (Figure 1 and 2) | GGCCCAGACTGAGCACGTGA |
| TNF.1 | TNF | Initial validation (Figure 1 and 2) | TGAAAGCATGATCCGGGACG |
| IL6.1 | IL6 | Initial validation (Figure 1 and 2) | TGAAAGCAGCAAAGAGGCAC |
| MAP2K1.1 | MAP2K1 | Initial validation (Figure 1 and 2) | GAAAGAAAGCTCCAGGTCTG |
| DNMT1.1 | DNMT1 | Initial validation (Figure 1 and 2) | GATTCCTGGTGCCAGAAACA |
| VEGFA.1 | VEGFA | Initial validation (Figure 1 and 2) | GATGTCTGCAGGCCAGATGA |
| CD209.1 | CD209 | Initial validation (Figure 1 and 2) | GCCCTCCACTAGGGCAAGGGT |
| **HACE sgRNA sequences for multiplex editing** | | | |
| **Set 1** | | | |
| CD209-1 | CD209 | Initial validation (Figure 1 and 2) | GCCCTCCACTAGGGCAAGGGT |
| RUNX1-1 | RUNX1 | Initial validation (Figure 1 and 2) | ATGAAGCACTGTGGGTACGA |
| VEGFA-1 | VEGFA | Initial validation (Figure 1 and 2) | GATGTCTGCAGGCCAGATGA |
| **Set 2** | | | |
| DNMT1-1 | DNMT1 | Initial validation (Figure 1 and 2) | GATTCCTGGTGCCAGAAACA |
| HEK3-1 | HEK3 | Initial validation (Figure 1 and 2) | GGCCCAGACTGAGCACGTGA |
| IL6-1 | IL6 | Initial validation (Figure 1 and 2) | TGAAAGCAGCAAAGAGGCAC |
| **sgRNA used for MEK1 screen** | | | |
| MEK1i1 | MAP2K1 | MEK1i resistance screen (Figure 3) | GAAAGAAAGCTCCAGGTCTG |

| | | | |
|---|---|---|---|
| MEK1i2 | MAP2K1 | MEK1i resistance screen (Figure 3) | AAGCTCTTTAAGGTAGAGGG |
| MEK1i5 | MAP2K1 | MEK1i resistance screen (Figure 3) | ACACCCCCGTCCGCCATCAG |
| **sgRNA used for SF3B1 screen** | | | |
| SF3B1_1 | SF3B1 | SF3B1 missplicing screen (Figure 4) | ATAAGAAATTTAGAATTATC |
| SF3B1_2 | SF3B1 | SF3B1 missplicing screen (Figure 4) | AAGAAAGGACAGTCATGAGT |
| SF3B1_3 | SF3B1 | SF3B1 missplicing screen (Figure 4) | CCAGACTGGACTCAAACCTT |
| SF3B1_13 | SF3B1 | SF3B1 missplicing screen (Figure 4) | TGTTCACATTAAACAAAATT |
| SF3B1_5 | SF3B1 | SF3B1 missplicing screen (Figure 4) | ATTTATCTTCATTAAAGTTA |
| SF3B1_6 | SF3B1 | SF3B1 missplicing screen (Figure 4) | CCAGATTTGTTAATGTAAAC |
| SF3B1_7 | SF3B1 | SF3B1 missplicing screen (Figure 4) | TATATGTGCTTGATTATGAA |
| **sgRNA used for *CD69* enhancer screen** | | | |
| CD69_A | CD69 enhancer | CD69 enhancer screen(Figure 5) | CAAAGTGTGCAGAGAAGGTG |
| CD69_B | CD69 enhancer | CD69 enhancer screen(Figure 5) | TGTCTTAGGTCGGAAGTCTG |
| CD69_C | CD69 enhancer | CD69 enhancer screen(Figure 5) | GACTTGAAGAGGACAGAAGG |

**Supplementary Table 2. Table of primer sequence used for sequencing.**

| ID | Target | Sequence |
|---|---|---|
| **Amplicon Sequencing** | | |
| DC1200 | *HEK3* | TCGTCGGCAGCGTCAGATGTGTATAAGAGA<br>CAG GAGCTGCACATACTAGCCCC |
| DC1201 | *HEK3* | GTCTCGTGGGCTCGGAGATGTGTATAAGAG<br>ACAG AGGGAGCTTGGCATGAGAAA |
| DC642 | *TNF* | TCGTCGGCAGCGTCAGATGTGTATAAGAGA<br>CAGNNNNNN<br>GCAGAGGACCAGCTAAGAGG |
| DC643 | *TNF* | GTCTCGTGGGCTCGGAGATGTGTATAAGAG<br>ACAGCCCAG TCACTCCAAAGTGCAGCAGG |
| DC640 | *IL6* | TCGTCGGCAGCGTCAGATGTGTATAAGAGA<br>CAGNNNNNN TGCCAGGATGCCAATGAGTT |
| DC641 | *IL6* | GTCTCGTGGGCTCGGAGATGTGTATAAGAG<br>ACAGCCCAG CACTGCATGCAAGAGGGAGA |
| DC381 | *MEK1* | TCGTCGGCAGCGTCAGATGTGTATAAGAGA<br>CAGNNNNN TGGTGATAGTCATCCCGGGT |
| DC497 | *MEK1* | GTCTCGTGGGCTCGGAGATGTGTATAAGAG<br>ACAGCCCAG CGTCATCCTTCAGTTCTCCC |
| DC1198 | *DNMT1* | TCGTCGGCAGCGTCAGATGTGTATAAGAGA<br>CAG TGTTCCCCAGAGTGACTTTTCC |
| DC1199 | *DNMT1* | GTCTCGTGGGCTCGGAGATGTGTATAAGAG<br>ACAG TTCCACTCATACAGTGGTAGATTT |
| DC1206 | *VEGFA* | TCGTCGGCAGCGTCAGATGTGTATAAGAGA<br>CAG GGGTTTTGCCAGACTCCACA |
| DC1207 | *VEGFA* | GTCTCGTGGGCTCGGAGATGTGTATAAGAG<br>ACAG TTGGGACTGGAGTTGCTTCA |
| DC1208 | *CD209* | TCGTCGGCAGCGTCAGATGTGTATAAGAGA<br>CAG AACAGGAAGTTGGGTAGGGA |
| DC1209 | *CD209* | GTCTCGTGGGCTCGGAGATGTGTATAAGAG<br>ACAG GAGGACAGCAGCAGCTCAAA |
| DC317 | *RUNX1* | TCGTCGGCAGCGTCAGATGTGTATAAGAGA<br>CAGNNNNN<br>ACAAACAAGACAGGGAACTGG |
| DC318 | *RUNX1* | GTCTCGTGGGCTCGGAGATGTGTATAAGAG<br>ACAGCCCAG<br>CTAGAGGGGTGAGGCTGAAAC |

| | | |
|---|---|---|
| DC1238 | *HEK3* long range | AACTAAACAGTCCCACTCCATCC |
| DC1239 | *HEK3* long range | GTATCTTCATGCATTCTCCACGCC |
| DC895 | *TNF* long range | GGAGAAACAGAGACAGGCCC |
| DC896 | *TNF* long range | CCAGGTTTCGAAGTGGTGGT |
| DC891 | *IL6* long range | CCCACCGGGAACGAAAGAGA |
| DC892 | *IL6* long range | GTCTCCCATTAGACCACAAGCA |
| **MEK1 screen** | | |
| GW_73 | MEK1 exon 2 | TCGTCGGCAGCGTCAGATGTGTATAAGAGACAGNNNNNNNNNNTTGTGCTCCCCACTTTGGAA |
| GW_74 | MEK1 exon 2 | GTCTCGTGGGCTCGGAGATGTGTATAAGAGACAGNNNNNNNNNNCCTGTTAATCAAGGCAAACTCACC |
| GW_77 | MEK1 exon 3 | TCGTCGGCAGCGTCAGATGTGTATAAGAGACAGNNNNNNNNNNGACTATATCTTTCATCCCTTCCTCCC |
| GW_78 | MEK1 exon 3 | GTCTCGTGGGCTCGGAGATGTGTATAAGAGACAGNNNNNNNNNNCAACTCTTAAGGCCATTGCTCC |
| GW_81 | MEK1 exon 6 | TCGTCGGCAGCGTCAGATGTGTATAAGAGACAGNNNNNNNNNNCCCAATCTACCTGTGTCAGTTCC |
| GW_82 | MEK1 exon 6 | GTCTCGTGGGCTCGGAGATGTGTATAAGAGACAGNNNNNNNNNNCCTACCCAGCACAAGACTCTG |
| DC1247 | MEK1 CDS | GGAGTTGGAAGCGCGTTAC |
| DC1248 | MEK1 CDS | CAAAAGCGACATGGCAAACC |
| **SF3B1 screen** | | |
| DC1510 | SF3B1 exon 12 | TCGTCGGCAGCGTCAGATGTGTATAAGAGACAG TCTGCTCTTTTCCCAGGCT |
| DC1511 | SF3B1 exon 12 | GTCTCGTGGGCTCGGAGATGTGTATAAGAGACAG ATTAAGGAGAACAAACCTTATGCAC |
| DC1512 | SF3B1 exon 13 | TCGTCGGCAGCGTCAGATGTGTATAAGAGACAG CTCGTGGTCATTGAACCGCT |
| DC1513 | SF3B1 exon 13 | GTCTCGTGGGCTCGGAGATGTGTATAAGAGACAG AGAAAGGACAGTCATGAGTTGGT |

| | | |
|---|---|---|
| DC1354 | SF3B1 exon 14 | TCGTCGGCAGCGTCAGATGTGTATAAGAGACAG TTCTTTGTTTACATTTTAGGCTGCT |
| DC1355 | SF3B1 exon 14 | GTCTCGTGGGCTCGGAGATGTGTATAAGAGACAG CTTCTAAGATGTGGCAAGATGGC |
| DC1356 | SF3B1 exon 15 | TCGTCGGCAGCGTCAGATGTGTATAAGAGACAG TTGTGTAACTTAGGTAATGTTGGGGC |
| DC1357 | SF3B1 exon 15 | GTCTCGTGGGCTCGGAGATGTGTATAAGAGACAG ACTTTAATGAAGATAAATCAAAAGG |
| DC1514 | SF3B1 exon 16 | TCGTCGGCAGCGTCAGATGTGTATAAGAGACAG GGCTGCTTTCTTGAAGGCTAT |
| DC1515 | SF3B1 exon 16 | GTCTCGTGGGCTCGGAGATGTGTATAAGAGACAG CTGTTAGAACCATGAAACATATCCA |
| DC1516 | SF3B1 exon 17 | TCGTCGGCAGCGTCAGATGTGTATAAGAGACAG ACAGTGTTGTGGGACAGATGG |
| DC1517 | SF3B1 exon 17 | GTCTCGTGGGCTCGGAGATGTGTATAAGAGACAG ACATGCATTCAAGTTGACTAAAGAA |
| DC1255 | SF3B1 CDS | CGTGAATTTGGAGCTGGTCC |
| DC1254 | SF3B1 CDS | ACGAATAGCCTTTTTGTGGGC |
| ***CD69* Enhancer Screen** | | |
| CD69 N3F | CD69 Enhancer | TACACGACGCTCTTCCGATCTAATCCAGGGTGAGACGTCAG |
| CD69 N3R | CD69 Enhancer | GGAGTTCAGACGTGTGCTCTTCCGATCTTGAGAGAGGATGATGTATTCTAAAT |
| CD69 N4F | CD69 Enhancer | TACACGACGCTCTTCCGATCTTCAGTGGGTGAATTAGGTTTCTGA |
| CD69 N4R | CD69 Enhancer | GGAGTTCAGACGTGTGCTCTTCCGATCTTGTGTTTCAGATGTTCCATTGAAG |
| CD69 BE-F | CD69 Enhancer | TACACGACGCTCTTCCGATCTTGGTGAGACGTCAGAAAGGAAGT |
| CD69 BE-R | CD69 Enhancer | GGAGTTCAGACGTGTGCTCTTCCGATCTAATTCACCCACTGAAAGGAAA |

**Supplementary Table 3. Table of helicase constructs tested.**

| Helicase | Species | Directionality |
|---|---|---|
| BLM[9] | Human | 3’ to 5’ |
| Ns3h[10] | Hepatitis C Virus | 3’ to 5’ |
| PcrA[11] | *Bacillus* | 3’ to 5’ |
| PcrA M6[12] | G. *stearothermophilus*, engineered | 3’ to 5’ |
| TraI[13] | E. *coli* | 5’ to 3’ |
| UvrD[14] | E. *coli* | 3’ to 5’ |

**Supplementary Table 4. Table of helicase sequences.**

| Helicase | Sequence |
|---|---|
| BLM | ATGGCTGCTGTTCCTCAAAATAATCTACAGGAGCAACTAGAACGTCACTC<br>AGCCAGAACACTTAATAATAAATTAAGTCTTTCAAAACCAAAATTTTCAG<br>GTTTCACTTTTAAAAAGAAAACATCTTCAGATAACAATGTATCTGTAACTA<br>ATGTGTCAGTAGCAAAAACACCTGTATTAAGAAATAAAGATGTTAATGTTA<br>CCGAAGACTTTTCCTTCAGTGAACCTCTACCCAACACCACAAATCAGCAA<br>AGGGTCAAGGACTTCTTTAAAAATGCTCCAGCAGGACAGGAAACACAGA<br>GAGGTGGATCAAAATCATTATTGCCAGATTTCTTGCAGACTCCGAAGGAA<br>GTTGTATGCACTACCCAAAACACACCAACTGTAAAGAAATCCCGGGATAC<br>TGCTCTCAAGAAATTAGAATTTAGTTCTTCACCAGATTCTTTAAGTACCAT<br>CAATGATTGGGATGATATGGATGACTTTGATACTTCTGAGACTTCAAAATC<br>ATTTGTTACACCACCCCAAAGTCACTTTGTAAGAGTAAGCACTGCTCAGA<br>AATCAAAAAAGGGTAAGAGAAACTTTTTAAAGCACAGCTTTATACAACA<br>AACACAGTAAAGACTGACTTGCCTCCACCCTCCTCTGAAAGCGAGCAAA<br>TAGATTTGACTGAGGAACAGAAGGATGACTCAGAATGGTTAAGCAGCGA<br>TGTGATTTGCATCGATGATGGCCCCATTGCTGAAGTGCATATAAATGAAGA<br>TGCTCAGGAAAGTGACTCTCTGAAAACTCATTTGGAAGATGAAAGAGAT<br>AATAGCGAAAAGAAGAAGAATTTGGAAGAAGCTGAATTACATTCAACTG<br>AGAAAGTTCCATGTATTGAATTTGATGATGATGATTATGATACGGATTTTGT<br>TCCACCTTCTCCAGAAGAAATTATTTCTGCTTCTTCTTCCTCTTCAAAATG<br>CCTTAGTACGTTAAAGGACCTTGACACCTCTGACAGAAAAGAGGATGTTC<br>TTAGCACATCAAAAGATCTTTTGTCAAAACCTGAGAAAATGAGTATGCAG<br>GAGCTGAATCCAGAAACCAGCACAGACTGTGACGCTAGACAGATAAGTT<br>TACAGCAGCAGCTTATTCATGTGATGGAGCACATCTGTAAATTAATTGATA<br>CTATTCCTGATGATAAACTGAAACTTTTGGATTGTGGGAACGAACTGCTTC<br>AGCAGCGGAACATAAGAAGGAAACTTCTAACGGAAGTAGATTTTAATAA<br>AAGTGATGCCAGTCTTCTTGGCTCATTGTGGAGATACAGGCCTGATTCAC<br>TTGATGGCCCTATGGAGGGTGATTCCTGCCCTACAGGGAATTCTATGAAG<br>GAGTTAAATTTTTCACACCTTCCCTCAAATTCTGTTTCTCCTGGGGACTGT<br>TTACTGACTACCACCCTAGGAAAGACAGGATTCTCTGCCACCAGGAAGA<br>ATCTTTTTGAAAGGCCTTTATTCAATACCCATTTACAGAAGTCCTTTGTAA<br>GTAGCAACTGGGCTGAAACACCAAGACTAGGAAAAAAAAATGAAAGCT<br>CTTATTTCCCAGGAAATGTTCTCACAAGCACTGCTGTGAAAGATCAGAAT<br>AAACATACTGCTTCAATAAATGACTTAGAAAGAGAAACCCAACCTTCCTA<br>TGATATTGATAATTTTGACATAGATGACTTTGATGATGATGATGACTGGGA<br>AGACATAATGCATAATTTAGCAGCCAGCAAATCTTCCACAGCTGCCTATCA<br>ACCCATCAAGGAAGGTCGGCCAATTAAATCAGTATCAGAAAGACTTTCCT<br>CAGCCAAGACAGACTGTCTTCCAGTGTCATCTACTGCTCAAAATATAAAC<br>TTCTCAGAGTCAATTCAGAATTATACTGACAAGTCAGCACAAAATTTAGC<br>ATCCAGAAATCTGAAACATGAGCGTTTCCAAAGTCTTAGTTTTCCTCATAC<br>AAAGGAAATGATGAAGATTTTTCATAAAAAATTTGGCCTGCATAATTTTAG<br>AACTAATCAGCTAGAGGCGATCAATGCTGCACTGCTTGGTGAAGACTGTT<br>TTATCCTGATGCCGACTGGAGGTGGTAAGAGTTTGTGTTACCAGCTCCCT<br>GCCTGTGTTTCTCCTGGGGTCACTGTTGTCATTTCTCCCTTGAGATCACTT |

| | |
|---|---|
| | ATCGTAGATCAAGTCCAAAAGCTGACTTCCTTGGATATTCCAGCTACATAT<br>CTGACAGGTGATAAGACTGACTCAGAAGCTACAAATATTTACCTCCAGTT<br>ATCAAAAAAAGACCCAATCATAAAACTCCTATATGTCACTCCAGAAAAGA<br>TCTGTGCAAGTAACAGACTCATTTCTACTCTGGAGAATCTCTATGAGAGG<br>AAGCTCTTGGCACGTTTTGTTATTGATGAAGCACATTGTGTCAGTCAGTG<br>GGGACATGATTTTCGTCAAGATTACAAAAGAATGAATATGCTTCGCCAGA<br>AGTTTCCTTCTGTTCCGGTGATGGCTCTTACGGCCACAGCTAATCCCAGG<br>GTACAGAAGGACATCCTGACTCAGCTGAAGATTCTCAGACCTCAGGTGTT<br>TAGCATGAGCTTTAACAGACATAATCTGAAATACTATGTATTACCGAAAAA<br>GCCTAAAAAGGTGGCATTTGATTGCCTAGAATGGATCAGAAAGCACCACC<br>CATATGATTCAGGGATAATTTACTGCCTCTCCAGGCGAGAATGTGACACCA<br>TGGCTGACACGTTACAGAGAGATGGGCTCGCTGCTCTTGCTTACCATGCT<br>GGCCTCAGTGATTCTGCCAGAGATGAAGTGCAGCAGAAGTGGATTAATCA<br>GGATGGCTGTCAGGTTATCTGTGCTACAATTGCATTTGGAATGGGGATTGA<br>CAAACCGGACGTGCGATTTGTGATTCATGCATCTCTCCCTAAATCTGTGGA<br>GGGTTACTACCAAGAATCTGGCAGAGCTGGAAGAGATGGGGAAATATCTC<br>ACTGCCTGCTTTTCTATACCTATCATGATGTGACCAGACTGAAAAGACTTA<br>TAATGATGGAAAAAGATGGAAACCATCATACAAGAGAAACTCACTTCAAT<br>AATTTGTATAGCATGGTACATTACTGTGAAAATATAACAGAATGCAGGAGA<br>ATACAGCTTTTGGCCTACTTTGGTGAAAATGGATTTAATCCTGATTTTTGTA<br>AGAAACACCCAGATGTTTCTTGTGATAATTGCTGTAAAACAAAGGATTAT<br>AAAACAAGAGATGTGACTGACGATGTGAAAAGTATTGTAAGATTTGTTCA<br>AGAACATAGTTCATCACAAGGAATGAGAAATATAAAACATGTAGGTCCTT<br>CTGGAAGATTTACTATGAATATGCTGGTCGACATTTTCTTGGGGAGTAAGA<br>GTGCAAAAATCCAGTCAGGTATATTTGGAAAAGGATCTGCTTATTCACGA<br>CACAATGCCGAAAGACTTTTTAAAAAGCTGATACTTGACAAGATTTTGGA<br>TGAAGACTTATATATCAATGCCAATGACCAGGCGATCGCTTATGTGATGCT<br>CGGAAATAAAGCACAAACTGTACTAAATGGCAATTTAAAGGTAGACTTTA<br>TGGAAACAGAAAATTCCAGCAGTGTGAAAAAACAAAAAGCGTTAGTAGC<br>AAAAGTGTCTCAGAGGGAAGAGATGGTTAAAAAATGTCTTGGAGAACTT<br>ACAGAAGTCTGCAAATCTCTGGGGAAAGTTTTTGGTGTCCATTACTTCAA<br>TATTTTTAATACCGTCACTCTCAAGAAGCTTGCAGAATCTTTATCTTCTGAT<br>CCTGAGGTTTTGCTTCAAATTGATGGTGTTACTGAAGACAAACTGGAAAA<br>ATATGGTGCGGAAGTGATTTCAGTATTACAGAAATACTCTGAATGGACATC<br>GCCAGCTGAAGACAGTTCCCCAGGGATAAGCCTGTCCAGCAGCAGAGGC<br>CCCGGAAGAAGTGCCGCTGAGGAGCTTGACGAGGAAATACCCGTATCTT<br>CCCACTACTTTGCAAGTAAAACCAGAAATGAAAGGAAGAGGAAAAAGAT<br>GCCAGCCTCCCAAAGGTCTAAGAGGAGAAAAACTGCTTCCAGTGGTTCC<br>AAGGCAAAGGGGGGGTCTGCCACATGTAGAAAGATATCTTCCAAAACGA<br>AATCCTCCAGCATCATTGGATCCAGTTCAGCCTCACATACTTCTCAAGCGA<br>CATCAGGAGCCAATAGCAAATTGGGGATTATGGCTCCACCGAAGCCTATA<br>AATAGACCGTTTCTTAAGCCTTCATATGCATTCTCA |
| Ns3h | ATGGTGGACTTCATACCCGTTGAGTCTATGGAAACTACCATGCGGTCTCCG<br>GTCTTCACAGACAACTCAACCCCCCCGGCTGTACCGCAGACATTCCAAGT<br>GGCACATCTGCACGCTCCTACTGGCAGCGGCAAGAGCACCAAAGTGCCG<br>GCTGCGTATGCAGCCCAAGGGTACAAGGTGCTCGTCCTGAACCCGTCCGT |

| | |
|---|---|
| | TGCCGCCACCTTAGGGTTTGGGGCGTATATGTCCAAGGCACACGGTATCG<br>ACCCTAACATCAGAACTGGGGTAAGGACCATTACCACGGGCGGCTCCATT<br>ACGTACTCCACCTATGGCAAGTTCCTTGCCGACGGTGGCTGTTCTGGGGG<br>CGCCTATGACATCATAATATGTGATGAGTGCCACTCAACTGACTCGACTAC<br>CATCTTGGGCATCGGCACAGTCCTGGACCAAGCGGAGACGGCTGGAGCG<br>CGGCTCGTCGTGCTCGCCACCGCTACACCTCCGGGATCGGTTACCGTGCC<br>ACACCCCAATATCGAGGAAATAGGCCTGTCCAACAATGGAGAGATCCCCT<br>TCTATGGCAAAGCCATCCCCATTGAGGCCATCAAGGGGGGGAGGCATCTC<br>ATTTTCTGCCATTCCAAGAAGAAATGTGACGAGCTCGCCGCAAAGCTGAC<br>AGGCCTCGGACTGAACGCTGTAGCATATTACCGGGGCCTTGATGTGTCCG<br>TCATACCGCCTATCGGAGACGTCGTTGTCGTGGCAACAGACGCTCTAATG<br>ACGGGTTTCACCGGCGATTTTGACTCAGTGATCGACTGCAATACATGTGT<br>CACCCAGACAGTCGACTTCAGCTTGGATCCCACCTTCACCATTGAGACGA<br>CGACCGTGCCCCAAGACGCGGTGTCGCGCTCGCAACGGCGAGGTAGAAC<br>TGGCAGGGGTAGGAGTGGCATCTACAGGTTTGTGACTCCAGGAGAACGG<br>CCCTCGGGCATGTTCGATTCTTCGGTCCTGTGTGAGTGCTATGACGCGGG<br>CTGTGCTTGGTATGAGCTCACGCCCGCTGAGACCTCGGTTAGGTTGCGGG<br>CTTACCTAAATACACCAGGGTTGCCCGTCTGCCAGGACCATCTGGAGTTC<br>TGGGAGAGCGTCTTCACAGGCCTCACCCACATAGATGCCCACTTCCTGTC<br>CCAGACTAAACAGGCAGGAGACAACTTTCCTTACCTGGTGGCATATCAAG<br>CTACAGTGTGCGCCAGGGCTCAAGCTCCACCTCCATCGTGGGACCAAATG<br>TGGAAGTGTCTCATACGGCTGAAACCTACACTGCACGGGCCAACACCCC<br>TGCTGTATAGGCTAGGAGCCGTCCAAAATGAGGTCATCCTCACACACCCC<br>ATAACTAAATACATCATGGCATGCATGTCGGCTGACCTGGAGGTCGTCACT |
| PcrA | ATGAATGCCCTGCTGAACCATATGAATACAGAACAATCCGAGGCGGTAAA<br>GACCACAGAAGGCCCCTTGTTGATCATGGCGGGGGCTGGGAGTGGTAAA<br>ACGAGGGTCCTTACTCACCGAATAGCGTACTTGCTTGACGAAAAGGACGT<br>GAGTCCATATAACGTGCTTGCCATTACCTTCACAAACAAGGCTGCTAGAG<br>AGATGAAGGAAAGAGTCCAAAAACTTGTCGGTGACCAAGCGGAGGTCAT<br>TTGGATGTCTACCTTCCATTCTATGTGCGTTCGCATACTTCGGCGAGACGC<br>GGATAGGATTGGGATCGAACGGAACTTCACGATAATAGATCCTACAGATC<br>AAAAGTCTGTAATAAAAGATGTTCTCAAAAATGAGAATATAGATAGCAAA<br>AAATTTGAACCCCGAATGTTCATAGGTGCCATATCAAACTTGAAGAACGA<br>ACTCAAAACACCTGCGGACGCACAAAAGGAAGCAACAGACTACCACAG<br>TCAGATGGTCGCAACTGTTTATTCCGGCTACCAACGACAGCTGAGTCGGA<br>ATGAAGCACTGGATTTTGATGATCTGATCATGACTACTATTAACCTTTTTGA<br>AAGAGTACCGGAAGTGTTGGAATATTACCAAAACAAATTTCAATATATCC<br>ACGTTGATGAATACCAAGATACTAATAAGGCACAGTATACATTGGTAAAGC<br>TGCTGGCGTCAAAGTTTAAAAATCTTTGCGTGGTCGGGGATAGTGACCAG<br>AGCATATACGGTTGGCGCGGCGCCGACATACAGAATATCTTGTCCTTCGA<br>GAAAGATTATCCTGAGGCGAATACAATCTTCCTTGAGCAGAATTATAGATC<br>TACAAAAACTATTTTGAACGCGGCTAACGAAGTAATAAAAAATAATAGTG<br>AGCGAAAGCCTAAAGGTCTGTGGACAGCTAACACAAATGGTGAAAAGAT<br>TCATTACTACGAAGCAATGACTGAACGAGACGAAGCGGAGTTCGTCATCC<br>GGGAAATAATGAAACACCAACGCAACGGCAAGAAATACCAAGACATGGC<br>AATTCTGTACAGGACCAATGCGCAATCCAGAGTTCTCGAAGAAACCTTTA |

| | |
|---|---|
| | TGAAGAGCAATATGCCATACACGATGGTTGGAGGCCAAAAATTCTATGAT<br>AGGAAAGAGATCAAAGACCTGCTGAGCTACCTCCGAATCATTGCCAACA<br>GTAACGACGACATCTCACTTCAACGGATTATTAACGTACCGAAACGCGGG<br>GTTGGACCCTCATCAGTTGAGAAAGTTCAAAACTATGCGTTGCAGAACA<br>ATATTTCCATGTTTGACGCTCTTGGAGAAGCTGATTTTATCGGCTTGTCAA<br>AGAAAGTAACCCAGGAGTGTCTTAACTTTTACGAACTGATACAAAGCCTG<br>ATAAAGGAACAGGAATTCCTTGAGATCCACGAGATCGTAGATGAAGTTCT<br>GCAAAAATCCGGCTATCGGGAAATGTTGGAAAGGGAGAACACGCTCGAA<br>AGTAGGTCAAGACTCGAGAACATAGATGAGTTCATGTCAGTGCCCAAAG<br>ACTATGAGGAGAATACGCCCCTTGAAGAGCAGTCATTGATCAATTTCCTTA<br>CTGACCTGTCACTCGTTGCCGATATTGACGAAGCTGACACTGAGAATGGG<br>GTAACATTGATGACGATGCACAGTGCTAAGGGATTGGAGTTTCCCATAGT<br>CTTCATCATGGGTATGGAGGAGTCCCTCTTTCCACACATTCGGGCAATCAA<br>ATCCGAGGATGATCATGAGATGCAAGAGGAGCGCAGAATCTGTTACGTTG<br>CGATTACACGAGCGGAGGAGGTTCTTTATATTACACACGCAACGTCTCGG<br>ATGCTCTTTGGACGCCCACAGTCAAACATGCCCTCTAGGTTCCTTAAGGA<br>AATACCCGAGAGCCTGCTGGAGAACCATAGCTCAGGTAAGCGGCAGACC<br>ATACAACCAAAAGCTAAGCCGTTCGCCAAACGCGGCTTCTCACAGCGCA<br>CGACTAGCACGAAGAAGCAGGTGCTCTCCAGCGATTGGAATGTTGGCGA<br>TAAGGTTATGCACAAAGCTTGGGGAGAGGGCATGGTTTCCAATGTGAATG<br>AGAAAAATGGATCCATAGAGCTCGACATCATCTTTAAGAGCCAGGGCCCA<br>AAGCGCCTGCTCGCTCAGTTCGCTCCAATCGAGAAAAAAGAAGAC |
| PcrA M6 | ATGAACTTCCTGTCTGAGCAACTGCTGGCACACCTGAACAAGGAGCAGC<br>AAGAAGCTGTGCGGACCACCGAGGGACCTCTGCTGATCATGGCCGGCGC<br>TGGAAGCGGAAAAACAAGAGTGCTGACACACCGCATCGCCTACCTGATG<br>GCTGAGAAGCACGTGGCCCCTTGGAACATCCTGGCCATTACCTTTACAAA<br>CAAGGCCGCTAGAGAGATGAGAGAAAGAGTGCAGAGCCTCCTGGGAGG<br>CGCCGCCGAGGACGTGTGGATCAGCACCTTCGCCAGCATGGCCGTGCGG<br>ATCCTGAGAAGAGATATCGACAGAATCGGCATCAACCGGAACTTCAGCAT<br>CCTGGACCCAACAGACCAGCTGAGCGTGATGAAAACCATCCTGAAAGAA<br>AAGAACATCGACCCCAAGAAATTCGAGCCTAGAACAATCCTGGGCACAA<br>TCAGCGCCGCCAAGAATGAACTGCTGCCTCCAGAACAGTTTGCGAAGCG<br>GGCCTCCACCTACTATGAGAAAGTGGTGTCTGACGTCTACCAGGAGTATC<br>AGCAGCGGCTGCTCAGGTGTCACAGCCTTGACTTCGATGATCTGATCATG<br>ACCACAATCCAGCTGTTTGACCGGGTCCCCGACGTGCTCCACTACTACCA<br>ATACAAGTTTCAGTACATCCACATCGATGAGTACCAGGACACAAACAGAG<br>CCCAATACACCCTGGTGAAAAAGCTGGCTGAGCGGTTCCAGAACATCGC<br>CGCCGTGGGCGACGCCGATCAGTCTATCTACAGATGGCGGGGCGCCGAC<br>ATCCAGAACATCCTGAGCTTCGAAAGAGATTACCCCAACGCCAAAGTGAT<br>CCTGCTGGAGCAAAATTACCGGAGCACGAAGCGCATCCTGCAGGCCGCA<br>AACGAGGTGATCGAGCACAACGTGAACAGAAAGCCTAAGCGGATCTGGA<br>CCGAGAATCCTGAGGGCAAGCCCATCCTGTACTACGAGGCCATGAACGA<br>AGCCGACGAGGCCCAGTTCGTGGCCGGCAGAATCAGAGAGGCCGTGGA<br>GCGCGGCGAGAGAAGATACCGAGACTTCGCCGTGCTGTACAGAACCAAT<br>GCCCAGTCCAGAGTCATGGAAGAGATGCTGCTGAAGGCCAACATCCCTT<br>ACCAGATAGTGGGCGGCGTGAAGTTCTACGACAGAAAGGAAATCAAGGA |

| | |
|---|---|
| | CATCCTGGCTTATCTGCGGGTCATCGCAAACCCAGACGACGACTGCAGCC<br>TGCTGAGAATTATCAACGTTCCTAAAAGAGGCATCGGAGCCTCTACCATC<br>GACAAGCTGGTCCGGTACGCCGCCGACCACGAGCTGAGCCTGTTCGAGG<br>CCCTGGGCGAGCTGGAGATGATCGGTCTTGGCGCCAAGGCCGCTGGGGC<br>CCTGGCCGCCTTCAGAAGCCAGCTGGAACAGTGGACACAGCTGCAGGA<br>GTACGTGAGCGTGACCGAACTGGTGGAAGAGGTGCTGGACAAGTCTGGC<br>TACCGGGAAATGCTGAAGGCCGAAAGAACCATCGAAGCCCAGAGCAGA<br>CTGGAGAACCTGGATGAGTTCCTGAGCGTGACCAAGCATTTCGAAAACG<br>TGTCCGACGACAAGAGCCTGATCGCCTTCCTGACCGATCTGGCTCTGATC<br>AGCGACCTGGACGAGCTGGATGGCACCGAGCAGGCCGCCGAGGGCGAC<br>GCCGTGATGCTGATGACCCTGCACGCCGCCAAGGGCCTGGAATTCCCCGT<br>GGTCTTTCTGATAGGCATGGAAGAAGGCATCTTCCCTCACAATAGAAGCC<br>TGGAGGATGACGATGAGATGGAAGAGGAGAGAAGACTGGCCTACGTGG<br>GCATCACCAGAGCTGAAGAGGAACTGGTGCTGACCAGCGCCCAGATGCG<br>GACCCTGTTCGGCAACATTCAGATGGACCCTCCTAGCCGGTTCCTTAACG<br>AGATCCCCGCCCATCTGCTTGAGACAGCCAGCAGACGGCAGGCCGGAGC<br>TTCCAGGCCTGCCGTGAGCAGACCCCAGGCCTCTGGCGCTGTGGGCAGC<br>TGGAAGGTGGGCGATCGGGCAAATCACAGAAAGTGGGGCATCGGAACC<br>GTGGTTTCCGTGCGGGGCGGCGGAGATGATCAGGAACTGGACATCGCTT<br>TTCCATCTCCTATCGGCATTAAGAGACTGCTAGCTAAATTCGCCCCTATCG<br>AGAAGGTG |
| TraI | ATGGCGAAGATCCACATGGTCCTTCAGGGTAAAGGTGGGGTCGGAAAAA<br>GCGCAATCGCGGCGATCATAGCCCAGTACAAGATGGACAAAGGTCAGAC<br>GCCGCTTTGTATAGATACAGATCCAGTCAATGCCACGTTTGAGGGATATAA<br>GGCCCTTAATGTACGACGCCTTAACATCATGGCTGGGGATGAGATCAACA<br>GCCGCAACTTTGATACACTGGTCGAGCTCATCGCGCCCACCAAAGATGAC<br>GTAGTAATCGACAACGGTGCCTCATCTTTTGTTCCTCTGTCACACTATCTC<br>ATATCAAATCAAGTACCGGCACTCCTCCAGGAGATGGGACATGAGCTCGT<br>GATACATACCGTGGTAACTGGCGGCCAGGCATTGCTTGACACTGTAAGTG<br>GGTTCGCCCAGCTTGCCAGCCAGTTTCCAGCTGAAGCTCTGTTTGTCGTG<br>TGGCTGAATCCGTACTGGGGTCCAATTGAACACGAAGGGAAGTCATTCG<br>AGCAAATGAAAGCTTACACTGCTAATAAGGCTAGGGTCTCAAGCATTATC<br>CAAATCCCAGCCCTTAAGGAGGAGACATACGGACGAGACTTCTCCGATAT<br>GCTGCAAGAACGGTTGACTTTCGACCAGGCCCTCGCAGACGAATCTTTG<br>ACTATAATGACCCGGCAGAGGCTTAAAATAGTGAGACGCGGCCTTTTCGA<br>ACAATTGGACGCCGCAGCCGTTCTG |
| UvrD | ATGGACGTATCTTACCTTTTGGATTCATTGAATGACAAACAGCGCGAAGC<br>TGTAGCTGCCCCAAGATCCAACCTCTTGGTGTTGGCGGGTGCTGGCTCAG<br>GGAAGACCCGAGTTCTTGTACACAGAATCGCGTGGCTTATGTCTGTTGAA<br>AATTGCAGCCCTTACTCTATAATGGCCGTCACGTTTACGAATAAGGCAGCT<br>GCGGAAATGCGCCATCGAATTGGGCAGCTTATGGGCACTTCACAAGGTGG<br>CATGTGGGTGGGAACTTTTCACGGGCTCGCTCATCGGCTGTTGCGCGCAC<br>ATCACATGGACGCGAACCTGCCTCAAGATTTTCAGATCCTCGATTCCGAA<br>GATCAATTGCGCCTTCTGAAAAGACTGATCAAGGCAATGAACCTGGATGA<br>AAAGCAATGGCCACCCCGACAGGCAATGTGGTACATAAACAGCCAAAAA |

| | |
|---|---|
| | GACGAGGGTCTTCGACCACATCATATCCAAAGTTATGGTAACCCTGTTGA<br>ACAAACATGGCAGAAGGTTTATCAGGCGTACCAGGAAGCCTGTGACCGA<br>GCGGGTCTTGTCGATTTTGCAGAACTGCTCCTCAGAGCACATGAGTTGTG<br>GCTCAATAAACCTCACATCCTTCAGCATTATAGGGAAAGATTCACGAATAT<br>ACTGGTTGATGAGTTTCAGGATACAAACAATATCCAATATGCTTGGATTAG<br>ACTTCTCGCAGGAGACACGGGTAAGGTGATGATCGTCGGTGACGATGAC<br>CAATCAATTTACGGATGGCGGGGTGCCCAGGTGGAAAACATACAAAGATT<br>TCTTAACGACTTCCCTGGGGCGGAAACGATTAGGTTGGAGCAGAATTATC<br>GGAGTACTTCTAACATACTCAGTGCAGCTAACGCACTCATCGAGAACAAC<br>AACGGCCGACTGGGAAAGAAGCTTTGGACAGATGGCGCTGACGGTGAG<br>CCGATATCACTGTATTGCGCGTTTAACGAGCTTGACGAGGCCCGCTTCGTC<br>GTCAATAGAATAAAAACCTGGCAAGATAATGGGGGGGCGCTCGCTGAGT<br>GCGCTATTTTGTATCGGAGTAACGCGCAAAGCCGGGTTTTGGAGGAAGCA<br>CTTCTTCAGGCATCTATGCCATACCGGATATATGGAGGAATGCGATTTTTTG<br>AACGCCAGGAGATCAAAGATGCGCTTAGTTATCTTCGACTTATTGCGAATA<br>GAAATGATGATGCGGCCTTTGAGCGGGTTGTCAATACACCCACACGCGGG<br>ATAGGTGACAGAACTCTGGATGTTGTCAGGCAAACATCTAGAGACCGGC<br>AACTCACCCTTTGGCAGGCATGTAGAGAACTGCTCCAGGAAAAAGCACT<br>GGCTGGCCGCGCGGCATCAGCACTGCAAAGGTTTATGGAGCTGATTGAC<br>GCTCTTGCACAGGAAACAGCGGACATGCCCTTGCATGTCCAAACGGATA<br>GAGTGATCAAAGATTCAGGTCTTCGCACGATGTATGAACAAGAAAAAGG<br>GGAGAAAGGGCAGACTAGGATAGAGAACCTCGAAGAATTGGTTACAGCT<br>ACCCGCCAGTTTTCATACAATGAGGAAGACGAGGACCTGATGCCACTCCA<br>AGCTTTCTTGTCCCACGCTGCGCTCGAAGCCGGCGAGGGTCAAGCTGATA<br>CCTGGCAGGATGCGGTACAACTGATGACCCTCCACTCAGCCAAGGGTCT<br>GGAATTCCCTCAAGTGTTTATCGTCGGGATGGAGGAGGGTATGTTCCCAT<br>CCCAGATGTCTTTGGACGAAGGAGGACGACTCGAGGAGGAGCGGCGAC<br>TCGCCTATGTTGGTGTAACCCGAGCGATGCAGAAACTCACGTTGACGTAT<br>GCAGAGACGCGCAGGTTGTATGGGAAGGAGGTGTACCACCGACCCTCTA<br>GGTTCATTGGCGAACTTCCTGAAGAATGTGTGGAAGAAGTACGCCTGCG<br>GGCTACGGTATCTAGGCCCGTTAGCCATCAACGGATGGGGACTCCAATGG<br>TGGAGAATGACTCAGGCTATAAGCTGGGCCAAAGGGTCCGCCACGCGAA<br>ATTTGGTGAGGGCACCATCGTAAACATGGAAGGAAGTGGTGAACATTCA<br>AGGCTTCAAGTAGCCTTTCAGGGACAAGGGATTAAGTGGCTTGTGGCCG<br>CATACGCGCGACTCGAGAGCGTG |

**Supplementary Table 5. Table of significant exome off-target sites.**

| Genomic bin | Sample |
|---|---|
| chr1:44100001-44200000 | AID |
| chr1:145300001-145400000 | AID-BLM-UGI |
| chr1:152100001-152200000 | AID |
| chr1:152200001-152300000 | AID,AID-Ns3h-UGI |
| chr1:152300001-152400000 | AID |
| chr10:128100001-128200000 | AID |
| chr11:1100001-1200000 | AID |
| chr11:1200001-1300000 | AID |
| chr11:62500001-62600000 | AID,AID-Ns3h-UGI |
| chr14:104900001-105000000 | AID |
| chr4:9200001-9300000 | AID |
| chr5:140800001-140900000 | AID |
| chr5:141300001-141400000 | AID |
| chr6:26100001-26200000 | AID-Ns3h-UGI,AID-PcrA M6-UGI |
| chr6:27100001-27200000 | AID-Ns3h-UGI,AID-PcrA M6-UGI |
| chr6:31000001-31100000 | AID |
| chr6:31800001-31900000 | AID-Ns3h-UGI,AID-PcrA-UGI |
| chr7:73100001-73200000 | AID-PcrA-UGI |
| chr7:101000001-101100000 | AID |
| chr8:143800001-143900000 | AID |
| chrX:115100001-115200000 | AID |

**Supplementary Table 6. Table of hit candidates identified in MEK1 resistance screen.**

| Drug | CDS base position | Reference Base | Alternate Base | Phenotype | Adjusted p-value |
|---|---|---|---|---|---|
| Selumetinib | 170 | A | T | K57M | 1.37E-45 |
| Selumetinib | 360 | G | A | Silent | 1.18E-30 |
| Selumetinib | 383 | G | A | G128D | 5.51E-56 |
| Selumetinib | 607 | G | A | E203K | 0 |
| Selumetinib | 681 | G | A | Silent | 6.66E-24 |

| | | | | | |
|---|---|---|---|---|---|
| Trametinib | 383 | G | A | G128D | 0.00E+00 |
| Trametinib | 605 | G | A | G202E | 0.00E+00 |
| Trametinib | 607 | G | A | E203K | 3.98E-190 |
| Trametinib | 630 | G | A | Silent | 0.00E+00 |
| Trametinib | 654 | C | T | Silent | 0.04652943 |

**Supplementary Table 7. Table of CDS sequence for MEK1 mutations.**

| Name | Sequence |
|---|---|
| MEK1 WT | ATGCCCAAGAAGAAGCCGACGCCCATCCAGCTGAACCCGGCCCCC<br>GACGGCTCTGCAGTTAACGGGACCAGCTCTGCGGAGACCAACTTG<br>GAGGCCTTGCAGAAGAAGCTGGAGGAGCTAGAGCTTGATGAGCAG<br>CAGCGAAAGCGCCTTGAGGCCTTTCTTACCCAGAAGCAGAAGGTG<br>GGAGAACTGAAGGATGACGACTTTGAGAAGATCAGTGAGCTGGGG<br>GCTGGCAATGGCGGTGTGGTGTTCAAGGTCTCCCACAAGCCTTCTG<br>GCCTGGTCATGGCCAGAAAGCTAATTCATCTGGAGATCAAACCCGC<br>AATCCGGAACCAGATCATAAGGGAGCTGCAGGTTCTGCATGAGTGC<br>AACTCTCCGTACATCGTGGGCTTCTATGGTGCGTTCTACAGCGATGG<br>CGAGATCAGTATCTGCATGGAGCACATGGATGGAGGTTCTCTGGATC<br>AAGTCCTGAAGAAAGCTGGAAGAATTCCTGAACAAATTTTAGGAA<br>AAGTTAGCATTGCTGTAATAAAAGGCCTGACATATCTGAGGGAGAA<br>GCACAAGATCATGCACAGAGATGTCAAGCCCTCCAACATCCTAGTC<br>AACTCCCGTGGGGAGATCAAGCTCTGTGACTTTGGGGTCAGCGGG<br>CAGCTCATCGACTCCATGGCCAACTCCTTCGTGGGCACAAGGTCCT<br>ACATGTCGCCAGAAAGACTCCAGGGGACTCATTACTCTGTGCAGTC<br>AGACATCTGGAGCATGGGACTGTCTCTGGTAGAGATGGCGGTTGGG<br>AGGTATCCCATCCCTCCTCCAGATGCCAAGGAGCTGGAGCTGATGT<br>TTGGGTGCCAGGTGGAAGGAGATGCGGCTGAGACCCCACCCAGGC<br>CAAGGACCCCCGGGAGGCCCCTTAGCTCATACGGAATGGACAGCC<br>GACCTCCCATGGCAATTTTTGAGTTGTTGGATTACATAGTCAACGAG<br>CCTCCTCCAAAACTGCCCAGTGGAGTGTTCAGTCTGGAATTTCAAG<br>ATTTTGTGAATAAATGCTTAATAAAAAACCCCGCAGAGAGAGCAGA<br>TTTGAAGCAACTCATGGTTCATGCTTTTATCAAGAGATCTGATGCTG<br>AGGAAGTGGATTTTGCAGGTTGGCTCTGCTCCACCATCGGCCTTAA<br>CCAGCCCAGCACACCAACCCATGCTGCTGGCGTCTAA |
| MEK1 G128D | ATGCCCAAGAAGAAGCCGACGCCCATCCAGCTGAACCCGGCCCCC<br>GACGGCTCTGCAGTTAACGGGACCAGCTCTGCGGAGACCAACTTG<br>GAGGCCTTGCAGAAGAAGCTGGAGGAGCTAGAGCTTGATGAGCAG<br>CAGCGAAAGCGCCTTGAGGCCTTTCTTACCCAGAAGCAGAAGGTG<br>GGAGAACTGAAGGATGACGACTTTGAGAAGATCAGTGAGCTGGGG<br>GCTGGCAATGGCGGTGTGGTGTTCAAGGTCTCCCACAAGCCTTCTG<br>GCCTGGTCATGGCCAGAAAGCTAATTCATCTGGAGATCAAACCCGC<br>AATCCGGAACCAGATCATAAGGGAGCTGCAGGTTCTGCATGAGTGC<br>AACTCTCCGTACATCGTGGACTTCTATGGTGCGTTCTACAGCGATGG<br>CGAGATCAGTATCTGCATGGAGCACATGGATGGAGGTTCTCTGGATC<br>AAGTCCTGAAGAAAGCTGGAAGAATTCCTGAACAAATTTTAGGAA<br>AAGTTAGCATTGCTGTAATAAAAGGCCTGACATATCTGAGGGAGAA<br>GCACAAGATCATGCACAGAGATGTCAAGCCCTCCAACATCCTAGTC<br>AACTCCCGTGGGGAGATCAAGCTCTGTGACTTTGGGGTCAGCGGG<br>CAGCTCATCGACTCCATGGCCAACTCCTTCGTGGGCACAAGGTCCT<br>ACATGTCGCCAGAAAGACTCCAGGGGACTCATTACTCTGTGCAGTC |

| | |
|---|---|
| | AGACATCTGGAGCATGGGACTGTCTCTGGTAGAGATGGCGGTTGGG<br>AGGTATCCCATCCCTCCTCCAGATGCCAAGGAGCTGGAGCTGATGT<br>TTGGGTGCCAGGTGGAAGGAGATGCGGCTGAGACCCCACCCAGGC<br>CAAGGACCCCCGGGAGGCCCCTTAGCTCATACGGAATGGACAGCC<br>GACCTCCCATGGCAATTTTTGAGTTGTTGGATTACATAGTCAACGAG<br>CCTCCTCCAAAACTGCCCAGTGGAGTGTTCAGTCTGGAATTTCAAG<br>ATTTTGTGAATAAATGCTTAATAAAAAACCCCGCAGAGAGAGCAGA<br>TTTGAAGCAACTCATGGTTCATGCTTTTATCAAGAGATCTGATGCTG<br>AGGAAGTGGATTTTGCAGGTTGGCTCTGCTCCACCATCGGCCTTAA<br>CCAGCCCAGCACACCAACCCATGCTGCTGGCGTCTAA |
| MEK1 G202E | ATGCCCAAGAAGAAGCCGACGCCCATCCAGCTGAACCCGGCCCCC<br>GACGGCTCTGCAGTTAACGGGACCAGCTCTGCGGAGACCAACTTG<br>GAGGCCTTGCAGAAGAAGCTGGAGGAGCTAGAGCTTGATGAGCAG<br>CAGCGAAAGCGCCTTGAGGCCTTTCTTACCCAGAAGCAGAAGGTG<br>GGAGAACTGAAGGATGACGACTTTGAGAAGATCAGTGAGCTGGGG<br>GCTGGCAATGGCGGTGTGGTGTTCAAGGTCTCCCACAAGCCTTCTG<br>GCCTGGTCATGGCCAGAAAGCTAATTCATCTGGAGATCAAACCCGC<br>AATCCGGAACCAGATCATAAGGGAGCTGCAGGTTCTGCATGAGTGC<br>AACTCTCCGTACATCGTGGGCTTCTATGGTGCGTTCTACAGCGATGG<br>CGAGATCAGTATCTGCATGGAGCACATGGATGGAGGTTCTCTGGATC<br>AAGTCCTGAAGAAAGCTGGAAGAATTCCTGAACAAATTTTAGGAA<br>AAGTTAGCATTGCTGTAATAAAAGGCCTGACATATCTGAGGGAGAA<br>GCACAAGATCATGCACAGAGATGTCAAGCCCTCCAACATCCTAGTC<br>AACTCCCGTGAGGAGATCAAGCTCTGTGACTTTGGGGTCAGCGGG<br>CAGCTCATCGACTCCATGGCCAACTCCTTCGTGGGCACAAGGTCCT<br>ACATGTCGCCAGAAAGACTCCAGGGGACTCATTACTCTGTGCAGTC<br>AGACATCTGGAGCATGGGACTGTCTCTGGTAGAGATGGCGGTTGGG<br>AGGTATCCCATCCCTCCTCCAGATGCCAAGGAGCTGGAGCTGATGT<br>TTGGGTGCCAGGTGGAAGGAGATGCGGCTGAGACCCCACCCAGGC<br>CAAGGACCCCCGGGAGGCCCCTTAGCTCATACGGAATGGACAGCC<br>GACCTCCCATGGCAATTTTTGAGTTGTTGGATTACATAGTCAACGAG<br>CCTCCTCCAAAACTGCCCAGTGGAGTGTTCAGTCTGGAATTTCAAG<br>ATTTTGTGAATAAATGCTTAATAAAAAACCCCGCAGAGAGAGCAGA<br>TTTGAAGCAACTCATGGTTCATGCTTTTATCAAGAGATCTGATGCTG<br>AGGAAGTGGATTTTGCAGGTTGGCTCTGCTCCACCATCGGCCTTAA<br>CCAGCCCAGCACACCAACCCATGCTGCTGGCGTCTAA |
| MEK1 E203K | ATGCCCAAGAAGAAGCCGACGCCCATCCAGCTGAACCCGGCCCCC<br>GACGGCTCTGCAGTTAACGGGACCAGCTCTGCGGAGACCAACTTG<br>GAGGCCTTGCAGAAGAAGCTGGAGGAGCTAGAGCTTGATGAGCAG<br>CAGCGAAAGCGCCTTGAGGCCTTTCTTACCCAGAAGCAGAAGGTG<br>GGAGAACTGAAGGATGACGACTTTGAGAAGATCAGTGAGCTGGGG<br>GCTGGCAATGGCGGTGTGGTGTTCAAGGTCTCCCACAAGCCTTCTG<br>GCCTGGTCATGGCCAGAAAGCTAATTCATCTGGAGATCAAACCCGC<br>AATCCGGAACCAGATCATAAGGGAGCTGCAGGTTCTGCATGAGTGC<br>AACTCTCCGTACATCGTGGGCTTCTATGGTGCGTTCTACAGCGATGG |

| | |
|---|---|
| | CGAGATCAGTATCTGCATGGAGCACATGGATGGAGGTTCTCTGGATC<br>AAGTCCTGAAGAAAGCTGGAAGAATTCCTGAACAAATTTTAGGAA<br>AAGTTAGCATTGCTGTAATAAAAGGCCTGACATATCTGAGGGAGAA<br>GCACAAGATCATGCACAGAGATGTCAAGCCCTCCAACATCCTAGTC<br>AACTCCCGTGGGAAGATCAAGCTCTGTGACTTTGGGGTCAGCGGG<br>CAGCTCATCGACTCCATGGCCAACTCCTTCGTGGGCACAAGGTCCT<br>ACATGTCGCCAGAAAGACTCCAGGGGACTCATTACTCTGTGCAGTC<br>AGACATCTGGAGCATGGGACTGTCTCTGGTAGAGATGGCGGTTGGG<br>AGGTATCCCATCCCTCCTCCAGATGCCAAGGAGCTGGAGCTGATGT<br>TTGGGTGCCAGGTGGAAGGAGATGCGGCTGAGACCCCACCCAGGC<br>CAAGGACCCCCGGGAGGCCCCTTAGCTCATACGGAATGGACAGCC<br>GACCTCCCATGGCAATTTTTGAGTTGTTGGATTACATAGTCAACGAG<br>CCTCCTCCAAAACTGCCCAGTGGAGTGTTCAGTCTGGAATTTCAAG<br>ATTTTGTGAATAAATGCTTAATAAAAAACCCCGCAGAGAGAGCAGA<br>TTTGAAGCAACTCATGGTTCATGCTTTTATCAAGAGATCTGATGCTG<br>AGGAAGTGGATTTTGCAGGTTGGCTCTGCTCCACCATCGGCCTTAA<br>CCAGCCCAGCACACCAACCCATGCTGCTGGCGTCTAA |

**Supplementary Table 8. Table of sequences for SF3B1 splicing minigene reporter.**

| | |
|---|---|
| DLST | **ATGGGTCGCTTTGACAGGGAGGTAGATATTGGAATTCCTGATGCTA**<br>**CAGGACGCTTAGAGATTCTTCAGATCCATACCAAGAACATGAAGCT**<br>**GGCAGATGATGTGGACCTGGAACAG**GTGAAGTGATGATGATGGCTGA<br>CCAGGCGTTACAGTGTCTCTAGGCAGTTGCTGGGAACTGGCTAGAGAC<br>ATAAGGTTAAGATGTGAGGAGATGGGTTTTGATTTCTGGACAGGGGAAA<br>GGAAGTAATCTGAGATTGAATCCAGGAAATGAAGCTTCGACACCGAGCT<br>CGGAAAGGGTTCTTATAACATACCTGCCAAAATGGGTTTTGTGGCTACTG<br>GAGGCTCCCCGCTAACAGGTAGCCTTGTAGCCTTTGATTGTCTTTTCAG**C**<br>**TGTTGGAGACACAGTTGCAGAAGATGAAGTGGTTTGTGAGATTGAA**<br>**ACTGACAAG**CAAGGCAGCGAGGGCAGAGGAAGTCTTCTAACATGCGG<br>TGACGTGGAGGAGAATCCCGGCCCTGTCAGTAAAGGTGAAGAACTGTT<br>CACCGGGGTGGTGCCCATCCTGGTCGAGCTGGACGGCGACGTAAACGG<br>CCACAAGTTCAGCGTGTCCGGCGAGGGCGAGGGCGATGCCACCTACGG<br>CAAGCTGACCCTGAAGTTCATCTGCACCACCGGCAAGCTGCCCGTGCCC<br>TGGCCCACCCTCGTGACCACCCTGACCTACGGCGTGCAGTGCTTCAGCC<br>GCTACCCCGACCACATGAAGCAGCACGACTTCTTCAAGTCCGCCATGCC<br>CGAAGGCTACGTCCAGGAGCGCACCATCTTCTTCAAGGACGACGGCAA<br>CTACAAGACCCGCGCCGAGGTGAAGTTCGAGGGCGACACCCTGGTGAA<br>CCGCATCGAGCTGAAGGGCATCGACTTCAAGGAGGACGGCAACATCCT<br>GGGGCACAAGCTGGAGTACAACTACAACAGCCACAACGTCTATATCATG<br>GCCGACAAGCAGAAGAACGGCATCAAGGTGAACTTCAAGATCCGCCAC<br>AACATCGAGGACGGCAGCGTGCAGCTCGCCGACCACTACCAGCAGAAC<br>ACCCCCATCGGCGACGGCCCCGTGCTGCTGCCCGACAACCACTACCTGA<br>GCACCCAGTCCGCCCTGAGCAAAGACCCCAACGAGAAGCGCGATCACA<br>TGGTCCTGCTGGAGTTCGTGACCGCCGCCGGGATCACTCTCGGTATGGA<br>TGAATTATATAAATAA |
| PLAC4 | **ATGGGTCGCTTTGACAGGGAGGTAGATATTGGAATTCCTGATGCTA**<br>**CAGGACGCTTAGAGATTCTTCAGATCCATACCAAGAACATGAAGCT**<br>**GGCAGATGATGTGGACCTGGAACAG**GTGAAGTGATGATGATGGCTGA<br>CCAGGCGTTACAGTGTCTCTAGGCAGTTGCTGGGAACTGGCTAGAGAC<br>ATAAGGTTAAGATGTGAGGAGATGGGTTTTGATTTCTGGACAGGGGAAA<br>GGAAGTAATCTGAGATTGAATCCAGGAAATGAAGCTTCGACACCGAGCT<br>CGCCTTTGACAACTGTCTTGTTACAAATGTTTGGTCCTTGCTTCTTTAAC<br>CTCTTAGTAAAGTTTGTGTATTCTAG**ATTACCACAGTTCCAGAGACAAT**<br>**GCTGGCACAAGGCTTCCAGCCCATCCTGTCACACTGACACGGAGA**<br>**ATGAAATCGTCCTGCCTCTGGGCTCCTTAGATCAG**CAAGGCAGCGA<br>GGGCAGAGGAAGTCTTCTAACATGCGGTGACGTGGAGGAGAATCCCGG<br>CCCTGTCAGTAAAGGTGAAGAACTGTTCACCGGGGTGGTGCCCATCCTG<br>GTCGAGCTGGACGGCGACGTAAACGGCCACAAGTTCAGCGTGTCCGGC<br>GAGGGCGAGGGCGATGCCACCTACGGCAAGCTGACCCTGAAGTTCATC<br>TGCACCACCGGCAAGCTGCCCGTGCCCTGGCCCACCCTCGTGACCACC<br>CTGACCTACGGCGTGCAGTGCTTCAGCCGCTACCCCGACCACATGAAGC<br>AGCACGACTTCTTCAAGTCCGCCATGCCCGAAGGCTACGTCCAGGAGC<br>GCACCATCTTCTTCAAGGACGACGGCAACTACAAGACCCGCGCCGAGG |

| | |
|---|---|
| | TGAAGTTCGAGGGCGACACCCTGGTGAACCGCATCGAGCTGAAGGGCA<br>TCGACTTCAAGGAGGACGGCAACATCCTGGGGCACAAGCTGGAGTACA<br>ACTACAACAGCCACAACGTCTATATCATGGCCGACAAGCAGAAGAACG<br>GCATCAAGGTGAACTTCAAGATCCGCCACAACATCGAGGACGGCAGCG<br>TGCAGCTCGCCGACCACTACCAGCAGAACACCCCCATCGGCGACGGCC<br>CCGTGCTGCTGCCCGACAACCACTACCTGAGCACCCAGTCCGCCCTGA<br>GCAAAGACCCCAACGAGAAGCGCGATCACATGGTCCTGCTGGAGTTCG<br>TGACCGCCGCCGGGATCACTCTCGGTATGGATGAATTATATAAATAA |

Upstream exon - **Orange highlight bold**
Upstream intron - Grey highlight
Downstream intron - Green highlight
Alternative 3' splice-inclusion - Red
Downstream Exon - **Bold**
T2A linker - Cyan highlight
EGFP - Yellow highlight

All other unannotated sequences are linker sequences used for cloning.

**Supplementary Table 9. Table of hit candidates identified for SF3B1 splicing screen.**

| Base Position | ref_base | alt_base | Phenotype | Mutation Type | log_fold_change | Mutation Type |
|---|---|---|---|---|---|---|
| 1627 | A | G | T543A | Missense | 1.38663646 | Non-clinical |
| 1629 | A | G | NA | Silent | 2.53851962 | Non-clinical |
| 1653 | A | G | NA | Silent | 1.93565474 | Non-clinical |
| 1666 | A | G | I556V | Missense | 1.45809111 | Non-clinical |
| 1677 | A | G | I559M | Missense | 1.7797861 | Non-clinical |
| 1682 | A | G | Y561C | Missense | 1.81357408 | Non-clinical |
| 1698 | A | G | NA | Silent | 1.69377311 | Non-clinical |
| 1709 | A | G | Y570C | Missense | 1.24133574 | Clinical |
| 1743 | A | G | NA | Silent | 1.15100255 | Non-clinical |
| 1751 | A | G | D584G | Missense | 1.00724764 | Non-clinical |
| 1757 | A | G | D586G | Missense | 1.72523491 | Clinical |
| 1760 | A | G | Y587C | Missense | 2.29587436 | Non-clinical |
| 1763 | A | G | Y588C | Missense | 1.7345498 | Non-clinical |
| 1768 | A | G | R590G | Missense | 1.17442273 | Non-clinical |
| 1789 | A | G | I597V | Missense | 1.10438786 | Non-clinical |
| 1795 | A | G | N599D | Missense | 1.93523264 | Non-clinical |
| 1807 | G | A | A603T | Missense | 1.01810563 | Non-clinical |
| 1822 | A | G | T608A | Missense | 1.17303987 | Non-clinical |
| 1837 | A | G | M613V | Missense | 1.14329741 | Non-clinical |
| 1849 | A | G | I617V | Missense | 2.46217362 | Non-clinical |
| 1851 | A | G | I617M | Missense | 1.63338506 | Non-clinical |
| 1855 | A | G | N619D | Missense | 1.44132924 | Non-clinical |
| 1868 | A | G | Y623C | Missense | 2.08956152 | Clinical |
| 1890 | A | G | NA | Silent | 1.92804782 | Non-clinical |
| 1932 | A | G | NA | Silent | 1.58256239 | Non-clinical |
| 1944 | A | G | NA | Silent | 1.92578678 | Non-clinical |
| 1946 | A | G | K649R | Missense | 1.2510653 | Non-clinical |
| 1985 | A | G | H662R | Missense | 1.30651587 | Clinical |
| 1987 | A | G | T663A | Missense | 1.26105039 | Clinical |
| 1993 | A | G | I665V | Missense | 2.07577179 | Non-clinical |

| | | | | | | |
|---|---|---|---|---|---|---|
| 1996 | A | G | K666E | Missense | 1.7787869 | Clinical |
| 2004 | A | G | NA | Silent | 2.20581348 | Non-clinical |
| 2006 | A | G | Q669R | Missense | 1.96012616 | Non-clinical |
| 2007 | A | G | NA | Silent | 1.05882839 | Non-clinical |
| 2009 | A | G | Q670R | Missense | 2.06464879 | Clinical |
| 2011 | A | G | I671V | Missense | 1.17983839 | Non-clinical |
| 2013 | A | G | I671M | Missense | 1.63347448 | Non-clinical |
| 2017 | A | G | I673V | Missense | 1.69952682 | Non-clinical |
| 2023 | A | G | M675V | Missense | 1.59048095 | Non-clinical |
| 2045 | A | G | H682R | Missense | 1.23817356 | Non-clinical |
| 2050 | A | G | R684G | Missense | 1.6647414 | Non-clinical |
| 2078 | G | A | G693D | Missense | 2.18806225 | Non-clinical |
| 2079 | T | G | NA | Silent | 1.94440594 | Non-clinical |
| 2083 | G | A | V695M | Missense | 1.8945899 | Non-clinical |
| 2085 | G | A | NA | Silent | 1.67273762 | Non-clinical |
| 2086 | G | A | D696N | Missense | 1.25908073 | Non-clinical |
| 2148 | A | G | NA | Silent | 1.49481957 | Non-clinical |
| 2156 | A | G | Y719C | Missense | 1.18694284 | Non-clinical |
| 2193 | A | G | NA | Silent | 2.051128 | Non-clinical |
| 2203 | A | G | I735V | Missense | 1.52109971 | Non-clinical |
| 2213 | A | G | H738R | Missense | 1.03392157 | Non-clinical |
| 2217 | A | G | NA | Silent | 1.18460958 | Non-clinical |
| 2220 | A | G | NA | Silent | 1.44002304 | Clinical |
| 2255 | A | G | Y752C | Missense | 1.05442597 | Clinical |
| 2260 | A | G | I754V | Missense | 1.64857116 | Non-clinical |
| 2269 | A | G | M757V | Missense | 2.08157125 | Non-clinical |
| 2282 | A | G | Y761C | Missense | 2.20515259 | Non-clinical |
| 2291 | A | G | Y764C | Missense | 1.47561896 | Non-clinical |
| 2294 | A | G | Y765C | Missense | 1.40710654 | Non-clinical |
| 2296 | A | G | T766A | Missense | 2.31058735 | Non-clinical |
| 2299 | A | G | R767G | Missense | 1.59807615 | Non-clinical |
| 2313 | A | G | NA | Silent | 1.8531005 | Non-clinical |

**Supplementary Table 10. Table of hit candidates identified for CD69 enhancer screen.**

| Base Position | ref_base | alt_base | Phenotype | Potential motif targeting | log2_fold_change |
|---|---|---|---|---|---|
| Chr12:9764879 | G | A | Reduced CD69 expression(validated) | Affecting ETS motif | -2.375045886 |
| Chr12:9764880 | G | A | Reduced CD69 expression(validated) | Affecting ETS motif | -2.639362938 |
| Chr12:9764930 | G | A | Reduced CD69 expression(screening) | Unclear | -3.28869167 |
| Chr12:9764976 | G | A | Reduced CD69 expression(screening) | Unclear | -2.203083513 |
| Chr12:9764996 | C | T | Reduced CD69 expression(validated) | Affecting RUNX motif | -3.085492338 |
| Chr12:9765002 | C | T | Reduced CD69 expression(screening) | Affecting RUNX motif | -2.710291572 |
| Chr12:9765072 | G | A | Reduced CD69 expression(screening) | Affecting ZSCAN4 motif | -2.963576938 |
| Chr12:9765102 | C | T | Reduced CD69 expression(screening) | Affecting STAT6 motif | -2.123218172 |
| Chr12:9765256 | G | A | Reduced CD69 expression(screening) | Affecting RelB motif | -2.421260686 |
| Below are the additional manually selected ones from Figure 5c | | | | | |
| Ch12:9764948 | C | T | Reduced CD69 expression(validated) | Affecting GATA motif | -1.148686331 |
| Ch12:9764995 | C | T | Reduced CD69 expression(validated) | Affecting RUNX motif | -1.553088638 |
| Ch12:9764998 | C | T | Reduced CD69 expression(validated) | Affecting RUNX motif | -1.442156962 |

**Supplementary Table 11. Table of base editor validation sgRNAs.**

| sgRNA name | Target gene/region | Experiment | sgRNA sequence | Target function |
|---|---|---|---|---|
| sgCtrl | AAVS1 | NA | GGGGCCACTAG GGACAGGAT | Safe harbor locus |
| sg383 | MEK1 | Validation for CDS base 383 (G>A) | TAGAAGCCCAC GATGTACGG | |
| sg607-1 | MEK1 | Validation for CDS base 607 (G>A) | CCCACGGGAGT TGACTAGGA | |
| sg607-2 | MEK1 | Validation for CDS base 607 (G>A) | CTTGATCTCCCC ACGGGAGT | |
| sg1682 | SF3B1 | Validation for CDS base 1849 (A>G) | CTGTACAAACT TGATGACTT | |
| sg1849 | SF3B1 | Validation for CDS base 1849 (A>G) | ATAGATAACATG GATGAGTA | |
| sg1868 | SF3B1 | Validation for CDS base 1868 (A>G) | GAGTATGTCCG TAACACAAC | |
| sg1996-1 | SF3B1 | Validation for CDS base 1996 (A>G) | ATTAAGATTGTA CAACAGAT | |
| sg1996-2 | SF3B1 | Validation for CDS base 1996 (A>G) | TGGTATTAAGAT TGTACAAC | |
| sgK700E-1 | SF3B1 | Validation positive control for K700E | CAGAAAGTTCG GACCATCAG | |
| sgK700E-2 | SF3B1 | Validation positive control for K700E | AGCAGAAAGTT CGGACCATC | |
| sg4948 | CD69 enhancer | Validation for chr12:9764948 C>T(Figure S6) | TCCTTTCTGAC GTCTCACCC | destroy GATA motif |
| sg4879 | CD69 enhancer | Validation for chr12:9764879/80 G>A(Figure S6) | TATCAGACAGC TGCAGCAGC | destroy potential IRF/STAT or IRF/ETS motif |
| sg4995 | CD69 enhancer | Validation for chr12:9764995/6/8 C>T(Figure 5 and S7) | GACCACAGACT TCCGACCTA | destroy RUNX motif, potential gain GATA motif |

**Supplementary Table 12. Table of prime editor validation sgRNAs.**

| pegRNA name | Target gene/region | Experiment | pegRNA sequence | Target function |
|---|---|---|---|---|
| peg1682-1 | SF3B1 | Validation for CDS base 1682 (A>G) | GAACAAACCTTATGCACATAGTTTTAGAGCTAGAAATAGCAAGTTAAAATAAGGCTAGTCCGTTATCAACTTGAAAAAGTGGCACCGAGTCGGTGCTgCAAACTTGATGACTTAGTTCGTCCATATGTGCATAAGGTTCCGAAAGATTGACGCGGTTCTATCTAGTTACGCGTTAAACCAACTAGAAATTTTTT | |
| peg1682-2 | SF3B1 | Validation for CDS base 1682 (A>G) | GTACTTGTGAAAGTTATTGATGTTTTAGAGCTAGAAATAGCAAGTTAAAATAAGGCTAGTCCGTTATCAACTTGAAAAAGTGGCACCGAGTCGGTGCGTTTGcACAGTATCCTATCAATAACTTTCACTCCCTTTATTGACGCGGTTCTATCTAGTTACGCGTTAAACCAACTAGAAATTTTTT | |
| peg1849 | SF3B1 | Validation for CDS base 1849 (A>G) | GAAGCTCTAGCTGTTGTGTTAGTTTTAGAGCTAGAAATAGCAAGTTAAAATAAGGCTAGTCCGTTATCAACTTGAAAAAGTGGCACCGAGTCGGTGCACCTGATgTAGATAACATGGATGAGTATGTCCGTAACACAACAGCTAGATAATAACATTGACGCGGTTCTATCTAGTTACGCGTTAAACCAACTAGAAATTTTTT | |
| peg1849.51-1 | SF3B1 | Validation for CDS base 1849 and 1951 (A>G) | GAAGCTCTAGCTGTTGTGTTAGTTTTAGAGCTAGAAATAGCAAGTTAAAATAAGGCTAGTCCGTTATCAACTTGAAAAAGTGGCACCGAGTCGGTGCGATgTgGATAACATGGATGAGTATGTCCGTAACACAACAGCTAGAGCGAAATATTTTGACGCGGTTCTATCTAGTTA | |

| | | | |
|---|---|---|---|
| | | | CGCGTTAAACCAACTAGAAATTTTTT |
| peg1849.51-2 | SF3B1 | Validation for CDS base 1849 and 1951 (A>G) | GAAGCTCTAGCTGTTGTGTTAGTTTTAGAGCTAGAAATAGCAAGTTAAAATAAGGCTAGTCCGTTATCAACTTGAAAAAGTGGCACCGAGTCGGTGCACCTGATgTgGATAACATGGATGAGTATGTCCGTAACACAACAGCTAGATAAATATTTTGACGCGGTTCTATCTAGTTACGCGTTAAACCAACTAGAAATTTTTT |
| peg1851-1 | SF3B1 | Validation for CDS base 1851 (A>G) | GAAGCTCTAGCTGTTGTGTTAGTTTTAGAGCTAGAAATAGCAAGTTAAAATAAGGCTAGTCCGTTATCAACTTGAAAAAGTGGCACCGAGTCGGTGCGATATgGATAACATGGATGAGTATGTCCGTAACACAACAGCTAGAGCACAGAACCTTGACGCGGTTCTATCTAGTTACGCGTTAAACCAACTAGAAATTTTTT |
| peg1851-2 | SF3B1 | Validation for CDS base 1851 (A>G) | GAAGCTCTAGCTGTTGTGTTAGTTTTAGAGCTAGAAATAGCAAGTTAAAATAAGGCTAGTCCGTTATCAACTTGAAAAAGTGGCACCGAGTCGGTGCACCTGATATgGATAACATGGATGAGTATGTCCGTAACACAACAGCTAGAAACAACACTTGACGCGGTTCTATCTAGTTACGCGTTAAACCAACTAGAAATTTTTT |
| peg1868 | SF3B1 | Validation for CDS base 1868 (A>G) | GAAGCTCTAGCTGTTGTGTTAGTTTTAGAGCTAGAAATAGCAAGTTAAAATAAGGCTAGTCCGTTATCAACTTGAAAAAGTGGCACCGAGTCGGTGCTGAGTgTGTCCGTAACACAACAGCTAGTAACAAATTTGACGCGGTTCTATCTAGTTACGCGTTAAACCAACTAGAAATTTTTT |

| | | | | |
|---|---|---|---|---|
| peg1996-1 | SF3B1 | Validation for CDS base 1996 (A>G) | GTCCTGGCAAGCGAGACACACGTTTTAGAGCTAGAAATAGCAAGTTAAAATAAGGCTAGTCCGTTATCAACTTGAAAAAGTGGCACCGAGTCGGTGCATCTTAAcACCAGTGTGTCTCGCTTGCAACCCTTCTTGACGCGGTTCTATCTAGTTACGCGTTAAACCAACTAGAAATTTTTT | |
| peg1996-2 | SF3B1 | Validation for CDS base 1996 (A>G) | GTCCTGGCAAGCGAGACACACGTTTTAGAGCTAGAAATAGCAAGTTAAAATAAGGCTAGTCCGTTATCAACTTGAAAAAGTGGCACCGAGTCGGTGCATCTcAAcACCAGTGTGTCTCGCTTGCAATTCTTCTTGACGCGGTTCTATCTAGTTACGCGTTAAACCAACTAGAAATTTTTT | |
| peg1996-3 | SF3B1 | Validation for CDS base 1996 (A>G) | GTGTGCAAAAGCAAGAAGTCCGTTTTAGAGCTAGAAATAGCAAGTTAAAATAAGGCTAGTCCGTTATCAACTTGAAAAAGTGGCACCGAGTCGGTGCATCTTAAcACCAGTGTGTCTCGCTTGCCAGGACTTCTTGCTTTTGCTTCTAAATTTGACGCGGTTCTATCTAGTTACGCGTTAAACCAACTAGAAATTTTTT | |
| pegWT | CD69 enhancer | Validation Control for chr12:9764995/6/8 C->T(Figure 5 and S7) | TGTCTTAGGTCGGAAGTCTGGTTTTAGAGCTAGAAATAGCAAGTTAAAATAAGGCTAGTCCGTTATCAACTTGAAAAAGTGGCACCGAGTCGGTGCAACAGGGACCACAGACTTCCGACCTAAG | none |
| peg4995 | CD69 enhancer | Validation for chr12:9764995 C->T(Figure 5 and S7) | TGTCTTAGGTCGGAAGTCTGGTTTTAGAGCTAGAAATAGCAAGTTAAAATAAGGCTAGTCCGTTATCAACTTGAAAAAGTGGCACCGAGTCGGTG | destroy RUNX motif |

| | | | | |
|---|---|---|---|---|
| | | | CAACAGGGATCACAGACTT CCGACCTAAG | |
| peg4996 | CD69 enhancer | Validation for chr12:9764996 C->T(Figure 5 and S7) | TGTCTTAGGTCGGAAGTCT GGTTTTAGAGCTAGAAATA GCAAGTTAAAATAAGGCTA GTCCGTTATCAACTTGAAA AAGTGGCACCGAGTCGGTG CAACAGGGACTACAGACTT CCGACCTAAG | destroy RUNX motif |
| peg4998 | CD69 enhancer | Validation for chr12:9764998 C->T(Figure 5 and S7) | TGTCTTAGGTCGGAAGTCT GGTTTTAGAGCTAGAAATA GCAAGTTAAAATAAGGCTA GTCCGTTATCAACTTGAAA AAGTGGCACCGAGTCGGTG CAACAGGGACCATAGACTT CCGACCTAAG | destroy RUNX motif |
| peg4995/6/8 | CD69 enhancer | Validation for chr12:9764995/6/8 C->T(Figure 5 and S7) | TGTCTTAGGTCGGAAGTCT GGTTTTAGAGCTAGAAATA GCAAGTTAAAATAAGGCTA GTCCGTTATCAACTTGAAA AAGTGGCACCGAGTCGGTG CAACAGGGATTATAGACTT CCGACCTAAG | destroy RUNX motif, gain GATA motif |

| **Nicking guides** | | |
|---|---|---|
| **ID** | **Paired pegRNA** | **Spacer sequence** |
| DC737 | peg1682-1 | GCTGATGTCTCCTACACTTG |
| DC738 | peg1682-2 | GAACAAACCTTATGCACATA |
| DC740 | peg1849 | TGCTGTTGTAGCCTCTGCCC |
| DC740 | peg1849.51-1 | TGCTGTTGTAGCCTCTGCCC |
| DC740 | peg1849.51-2 | TGCTGTTGTAGCCTCTGCCC |
| DC740 | peg1851-1 | TGCTGTTGTAGCCTCTGCCC |
| DC740 | peg1851-2 | TGCTGTTGTAGCCTCTGCCC |
| DC743 | peg1868 | GCTGTTGTAGCCTCTGCCCT |
| DC745 | peg1996-1 | ACTTCTAAGATGTGGCAAGA |
| DC745 | peg1996-2 | ACTTCTAAGATGTGGCAAGA |
| DC748 | peg1996-3 | GCAATAAAGAAGGAATGCCC |

**Supplementary Table 13. Comparison of HACE to state-of-the-art methods for nucleotide diversification.**

| **Method (ref)** | **Species** | **Target** | **Editing rate** | **Editing window** |
|---|---|---|---|---|
| TRACE[15] | Mammalian | Exogenous overexpression | ~2 $kbp^{-1}$ per 3 days | ~2000 bp per T7 promoter |
| TRIDENT[16] | Mammalian | Exogenous overexpression | ~2 $kbp^{-1}$ per 3 days | ~2000 bp per T7 promoter |
| CRISPR-X[17] | Mammalian | Endogenous | ~1 $kbp^{-1}$ per ~12 days | 10-50 bp per sgRNA |
| TAM[18] | Mammalian | Endogenous | ~1 kbp-1 per ~7 days | 10-50 bp per sgRNA |
| EvolvR[19] | E. coli | Endogenous | ~0.05 $kbp^{-1}$ per day | ~50 bp per sgRNA |
| OrthoRep[20] | Yeast | Exogenous overexpression | ~0.16 $kbp^{-1}$ per day | At least 5000 bp |
| VEGAS[21] | Mammalian | Exogenous overexpression | ~1 $kbp^{-1}$ per day | ~ 6000 bp |
| HACE (our method) | Human | Endogenous | ~2 $kbp^{-1}$ per 3 days | >1000 bp per sgRNA |
